# Ribosome-rescuer PELO catalyzes the oligomeric assembly of NLR family proteins *via* activating their ATPase

**DOI:** 10.1101/2022.10.05.511050

**Authors:** Xiurong Wu, Zhang-Hua Yang, Jianfeng Wu, Jiahuai Han

## Abstract

NOD-like receptors (NLRs) are pattern recognition receptors for diverse innate immune responses. Self-oligomerization after engagement with a ligand was a generally accepted model for the activation of each NLR. We report here that a catalyzer is required for the self-oligomerization. Protein pelota homolog (PELO, Dom34 in yeast), a well-known surveillance factor in translational quality control/ribosome rescue, interacts with all cytosolic NLRs and activates their ATPase. In the case of flagellin-initiated NLRC4 inflammasome activation, flagellin-bound NAIP5 recruits the first NLRC4 and then PELO is required for correctly assembling the rest of NLRC4s into the NLRC4 complex one by one by activating the NLRC4 ATPase. Stoichiometric and functional data revealed that PELO cannot be a structural constituent of NLRC4 inflammasome but a powerful catalyzer for its assembly. The catalytic role of PELO in the activation of cytosolic NLRs is unexpected and may break new ground for future studies of NLR family members.

## INTRODUCTION

NLR family members are evolution-derived intracellular PRRs for various pathogen-associated molecular patterns (PAMPs) and damage-associated molecular patterns (DAMPs). Numerous human diseases such as Blau syndrome, Crohn’s disease, early-onset sarcoidosis, cryopyrin-associated periodic fever syndrome or bare lymphocyte syndrome are linked to polymorphisms in certain NLR genes (Chen et al., 2009; Geddes et al., 2009). In addition to pattern recognition, NLRs participate in diverse biological processes ranging from antigen presentation, autophagy to embryonic development, and the functions of many NLRs are still unknown (Meunier and Broz, 2017). Structurally, NLRs share common C-terminal leucine-rich repeat (LRR) domains, a central NACHT domain and a variable N-terminal effector domain (Meunier and Broz, 2017). Most of the NLRs are expressed in cytoplasm, except CIITA and NLRX1, which are mainly expressed in nucleus and mitochondria, respectively (Moore et al., 2008; Raval et al., 2003). It is proposed that the crucial step in the activation of a given NLR lies in its oligomerization mediated by the NACHT domain, and the oligomeric complexes (e.g. the inflammasome or nodosome) function as platforms allowing the recruitment of adaptor(s) and effector(s) and signaling for inflammatory responses. The NACHT domain is characterized as the signal transduction ATPases with numerous domains (STAND) clade of the AAA+ ATPase superfamily and the oligomerization of NLRs is believed to be driven by hydrolysis of ATP (Danot et al., 2009; Koonin and Aravind, 2000; Lelpe et al., 2004). Auto-inhibition of NACHT ATPase in NLRs could be a mechanism for keeping NLRs at rest as a recent study showed that ligand binding to LRR of NLRP1 led to the gain of ATPase activity by the NACHT domain (Bauernfried et al., 2021). However, the mechanistic insight of how to activate ATPase in driving the oligomerization of NLRs is still largely unknown.

PELO is an evolutionarily conserved component of the ribosome-associated quality control machinery (Doma and Parker, 2006; Guydosh and Green, 2014; Pisareva et al., 2011; Shoemaker et al., 2010; Tsuboi et al., 2012). It was first identified as a gene involved in spermatogenesis in *Drosophila melanogaster* (Castrillon et al., 1993), and has been reported to participate in many physiological processes, such as epidermal homeostasis (Liakath-Ali et al., 2018), mitophagy (Wu et al., 2018), cerebellar neurogenesis (Terrey et al., 2021), and efficient virus replication in *Drosophila* (Wu et al., 2014). The involvement of PELO in immune responses has not yet been reported.

In this study, we found that PELO was associated with NLRP3 inflammasome and further revealed that it could interact with all cytosolic NLR family proteins. For the tested NLR proteins including inflammasomal and non-inflammasomal NLRs, PELO was required for their ligand-induced activation. Biochemical and cellular studies showed that PELO bound to NACHT and/or LRR domain(s) of NLRs and activated the ATPase in all NLRs *in vitro*; PELO promoted NLRC4 oligomerization when all required proteins were reconstituted in either cultured HEK293T cells or cell free mixtures; and the amount of PELO associating with NLRC4 excluded PELO as a structural constituent in matured inflammasome. Our data collectively indicated that PELO functioned as a catalyzer that activated the ATPase of given NLRs such as NLRC4 during the incorporation of NLRC4 into the extending NLRC4 oligomer. The catalysis is essential due to that activation of the ATPase is required for the NLR to incorporate into the complex correctly and in the right conformation. We also found that this newly emerged function of PELO is independent of its conventional role in ribosome rescue, and the possible competition for the usage of PELO protein could have functional influences.

## RESULTS

### PELO is an interacting protein of all cytosolic NLRs and controls the activation of NLRs

We have performed quantitative mass spectrometry-based analyses of the NLRP3 complex and detected PELO as an interacting protein of NLRP3 (**Figure S1A**, see Method section for detail (He et al., 2015)). When co-expressed in HEK293T cells, NLRP3 can be co-immunoprecipitated with PELO (**Figure 1A**). Incubation of purified recombinant His-tagged PELO protein with the cell lysates of HEK293T cells expressing Flag-NLRP3 led to the binding of PELO to NLRP3 as determined by immunoprecipitation (**Figure 1B**), confirming the direct interaction between PELO and NLRP3.

**Figure 1.**
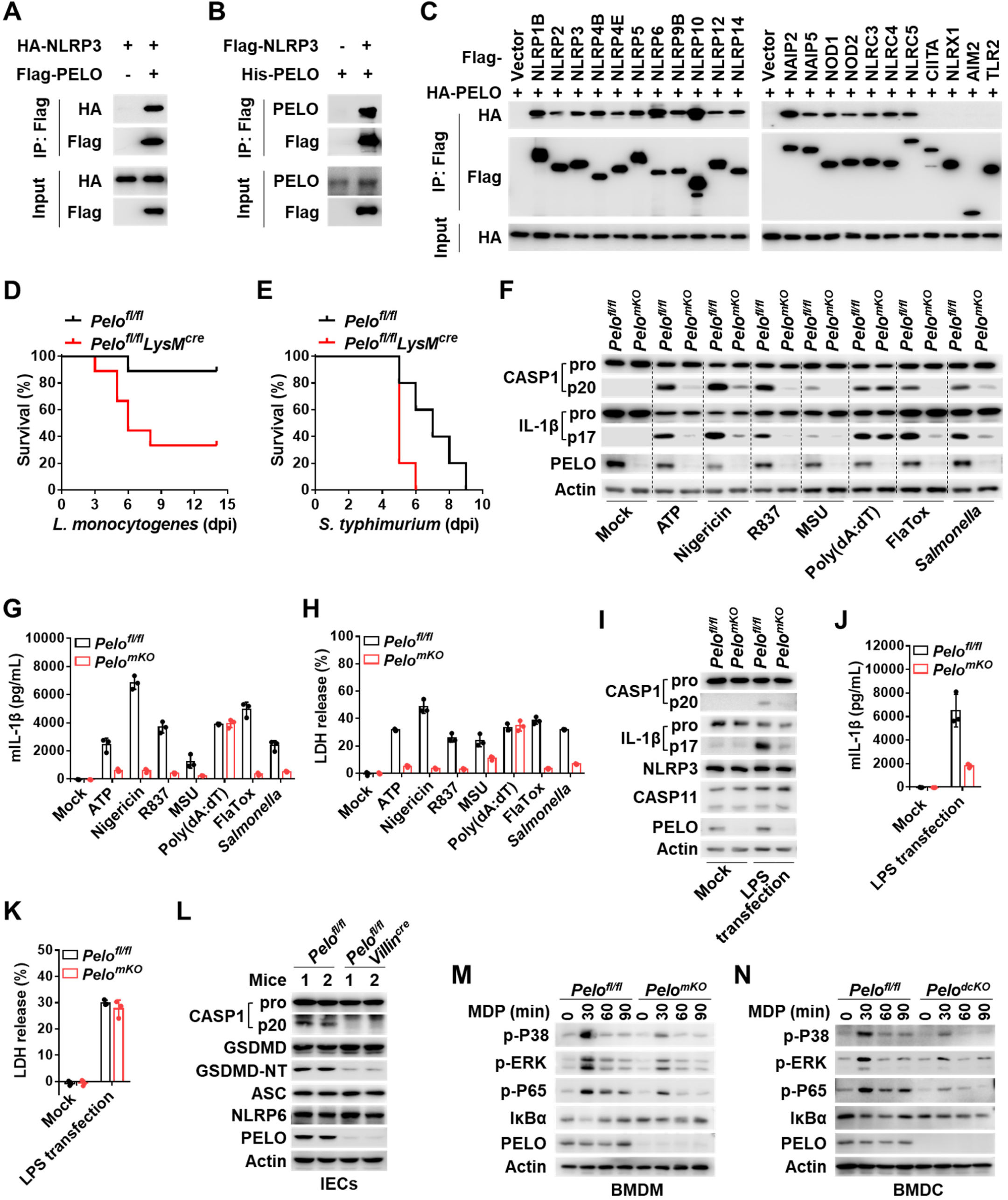
PELO interacts with all cytosolic NLRs and controls the activation of the NLRs tested. (**A**) HEK293T cells were transfected to overexpress HA-tagged NLRP3 or together with Flag-tagged PELO. The cell lysates were immunoprecipitated with anti-Flag antibodies. The cell lysates and immunoprecipitates were analyzed by immunoblotting as indicated. (**B**) Cell lysates from HEK293T cells transfected with or without Flag-tagged NLRP3 expression vector were incubated with recombinant His-tagged PELO protein, followed by immunoprecipitation with anti-Flag antibodies, and analyzed by immunoblotting. **(C**) Each of Flag-tagged NLRs was co-expressed with HA-tagged PELO in HEK293T cells. The cell lysates were immunoprecipitated with anti-Flag antibodies. The cell lysates and immunoprecipitates were analyzed by immunoblotting as indicated. (**D** and **E**) *Pelo^fl/fl^* and *Pelo^fl/fl^LysM^cre^* mice were infected by intraperitoneal injection of 1×10^6^ c.f.u. *L. monocytogenes* (**D**, n = 9 mice per genotype) or 1×10^3^ c.f.u. *S. typhimurium* (E, n = 9 mice per genotype), and survival was monitored daily. (**F**) Immunoblot analysis of the processed caspase-1 (CASP1) and IL-1β in the pooled cell extracts and supernatants from *Pelo^fl/fl^* and *Pelo^mKO^* BMDMs that were primed with LPS (100 ng/ml for 4 hours) and then treated with the indicated stimuli: 5 mM ATP (30 min), 5 μM nigericin (1 hour), 20 μg/ml R837 (1 hour), 200 μg/ml MSU (6 hours), 2 μg/ml poly(dA:dT) (2 hours), 2 μg/ml LFn-Flagellin together with 2 μg/ml PA proteins (1 hour), *Salmonella* (MOI=10, 1 hour). Mock represents BMDMs primed with LPS without further stimulation. (**G** and **H**) BMDMs were treated as in (**F**); IL-1β (**G**) and LDH (**H**) in culture supernatants were analyzed. n = 3 mice per genotype. (**I-K)** *Pelo^fl/fl^* and *Pelo^mKO^* BMDMs were primed with Pam3CSK4 (1 μg/ml) for 6 hours and then transfected with LPS (2 μg/ml). 12 hours post-transfection, the pooled cell extracts and supernatants were analyzed by immunoblotting (**I**); and the IL-1β secretion (**J**) or LDH release (**K**) from BMDMs were analyzed. n = 3 mice per genotype. (**L**) Immunoblot analysis of IEC lysates of *Pelo^fl/fl^* and *Pelo^fl/fl^Villin^cre^* mice. (**M** and **N**) Immunoblot analysis of the cell lysates from *Pelo^fl/fl^* and *Pelo^mKO^* BMDMs (**M**) or *Pelo^fl/fl^* and *Pelo^dcKO^* BMDCs (**N**) treated with L18-MDP at indicated time points. Data are represented as mean ± SD (**G**, **H, J** and **K**). All results are representative of at least two independent experiments. See also **Figure S1**, **S2** and **S3**.

To determine the domain(s) within NLRP3 that interact(s) with PELO, we expressed Flag-tagged wild-type (WT) NLRP3 or various NLRP3 truncations with HA-tagged PELO in HEK293T cells. PELO interacted with the NACHT and LRR domains but not the PYD domain of NLRP3 (**Figure S1B**). Since members of the NLR family contain the C-terminal LRR domain (except NLRP10) and central NACHT domain, we explored whether PELO could interact with other members of the NLR family using co-expression assay, and found that in addition to NLRP3, PELO interacted with all the other cytosolic NLRs, but not nuclear NLR CIITA or mitochondrial NLR NLRX1 (**Figure 1C**). The same as in NLRP3, NACHT and LRR domains in the other NLRs were responsible for the interaction with PELO (**Figure S1C-S1E**). To check whether PELO can bind to all kinds of PRRs, we tested whether PELO could co-immunoprecipitate with AIM2 or TLR2 by co-expressing them in HEK293T cells and did not detect PELO interaction with either of these two proteins (**Figure 1C**). Thus, PELO is a specific interacting protein of cytosolic NLRs.

To study PELO’s function in NLR-mediated immune responses, we generated *Pelo^fl/fl^* mice (**Figure S1F**), and bred *Pelo^fl/fl^* mice with *LysM^cre^* mice (*LysM* is also known as *Lyz2*) to delete *Pelo* from the myeloid compartment. *Pelo^fl/fl^LysM^cre^* mice were viable and did not display any obvious physical or behavioral abnormalities. The body weights of *Pelo^fl/fl^LysM^cre^* mice were comparable with those of *Pelo^fl/fl^* mice (**Figure S1G**). Bone marrow-derived macrophages (BMDMs) from *Pelo^fl/fl^LysM^cre^* mice (here named *Pelo^mKO^*) showed elimination of PELO protein (**Figure S1H**) but not the myeloid cell markers CD11b and F4/80 (**Figure S1I**). No defect in hematopoietic cell development was found in *Pelo^fl/fl^LysM^cre^* mice (**Figure S1J**). *Pelo* deletion in BMDMs had no detectable effect on the global protein expression in these cells under no-stress conditions (**Figure S1K** and **S1L**, and **Table S1**).

Functionally, NLRs could act as activators of NF-κB and mitogen-activated protein kinases (MAPKs) pathways, activators of inflammasomes, and regulators of proinflammatory responses (Babamale and Chen, 2021). As PELO could interact with all the cytosolic NLRs, we tested whether PELO plays roles in host defense against infections. Indeed, deletion of *Pelo* in myeloid cells rendered the mice much more susceptive to the death induced by either Gram-positive (*L. monocytogenes*) or Gram-negative (*S. typhimurium*) bacteria (**Figure 1D** and **1E**). Such defects in *Pelo* deficient mice should at least in part result from the impairments of NLRs’ activation by *Pelo* deficiency. We then initiated our study by examining the roles of PELO in the activation of several well-studied NLRs.

Some NLR proteins such as NLRP3 and NLRC4 are known for their roles in activating inflammasomes, the intracellular multimeric protein complexes that form in response to various exogenous microbial infections and endogenous danger signals (Broz and Dixit, 2016; Guo et al., 2015). Inflammasomes recruit the common adaptor protein ASC to activate inflammatory caspases such as caspase-1, which processes the proinflammatory cytokines interleukin 1β (IL-1β) and/or IL-18 for their maturation (Martinon et al., 2002), and cleaves gasdermin D (GSDMD) to generate the N-terminal fragment to induce pore formation, cytokine release and pyroptosis (Chen et al., 2016; Ding et al., 2016; He *et al*., 2015; Kayagaki et al., 2015; Liu et al., 2016; Shi et al., 2015). *Pelo^mKO^* and *Pelo^fl/fl^* (wild-type) BMDMs expressed comparable amounts of inflammasome-related proteins, including NLRC4, NEK7, ASC, GSDMD, caspase-1 and DDX3X (**Figure S2A**). The induction of NLRP3 and pro-IL-1β expression by lipopolysaccharide (LPS) was comparable between *Pelo^fl/fl^* and *Pelo^mKO^* cells (**Figure S2A**). Interestingly, activation of caspase-1 (**Figure 1F**), maturation and secretion of IL-1β (**Figure 1F** and **1G**), and pyroptosis (**Figure 1H** as indicated by LDH release) were impaired by *Pelo* deletion in LPS-primed BMDMs that were then treated with ATP, nigericin or MSU, the three potassium-efflux-dependent activators of NLRP3 inflammasome (Munoz-Planillo et al., 2013), as well as R837, a potassium-efflux-independent stimulus of NLRP3 inflammasome (Gross et al., 2016). Akin to NLRP3 inflammasome, activation of NLRC4 inflammasome by flagellin (Fla), which was achieved by using FlaTox comprised of recombinant *Legionella pneumophila* flagellin fused to the amino-terminal domain of *Bacillus anthracis* lethal factor and anthrax protective antigen (PA) (Kofoed and Vance, 2011; Zhao et al., 2011), or by *Salmonella enterica serovar Typhimurium* (*Salmonella*), was also impaired in *Pelo^mKO^*BMDMs (**Figure 1F-1H**). In contrast, activation of caspase-1, production of IL-1β and pyroptosis in response to poly(dA:dT), which activates the AIM2 inflammasome, were not affected in *Pelo^mKO^*BMDMs (**Figure 1F-1H**). This is consistent with that PELO could interact with NLRP3 and NLRC4 but not AIM2 (**Figure 1C**).

In the case of cytosolic LPS-induced noncanonical inflammasome activation, NLRP3-dependent caspase-1 activation and IL-1β production required PELO (**Figure 1I** and **1J**), but caspase-11-dependent pyroptosis did not (**Figure 1K**), consistent with the reported role of NLRP3 in the noncanonical inflammasome (Hagar et al., 2013; Kayagaki et al., 2011; Kayagaki et al., 2013). As expected, cytosolic LPS-induced TNFα production was not affected by *Pelo* deletion (**Figure S2B**). The induction of apoptosis by TNFα plus smac mimetic (SM164) or staurosporine (**Figure S2C** and **S2D**), necroptosis by TNF+SM164+zVAD or LPS+zVAD (**Figure S2E** and **S2F**), and ferroptosis by RSL3 (**Figure S2G**), was similar between *Pelo^fl/fl^* and *Pelo^mKO^* BMDMs. Altogether, these results demonstrated a specific requirement of PELO in NLRP3 and NLRC4 inflammasome activation.

NLRP6 is another inflammasomal NLR, which is expressed predominantly in mucosal tissues that are constantly exposed to microbial components (Venuprasad and Theiss, 2021). Recent studies have found that wild-type mice, but not *Nlrp6^-/-^* mice, showed robust caspase-1 and GSDMD processing in intestinal epithelial cells (IECs) at a steady state (Levy et al., 2015; Shen et al., 2021). To test the role of PELO in this NLRP6 inflammasome activation, we examined IECs from *Pelo^fl/fl^* and *Pelo^fl/fl^Villin^cre^* mice and found that absence of PELO in IECs also dramatically reduced processing of caspase-1 and GSDMD (**Figure 1L**), suggesting an essential role of PELO in NLRP6-mediated activation of caspase-1 and GSDMD.

NOD2 is a well-studied non-inflammasomal NLR that activates NF-κB and MAPKs after its recognition of bacterial peptidoglycan derivatives muramyl dipeptide (MDP) (Girardin et al., 2003; Inohara et al., 2003). We found that MDP induced NF-κB and MAPKs activations were effectively attenuated by *Pelo* deficiency in BMDMs (**Figure 1M**) and in bone marrow-derived dendritic cells (BMDCs) (here named *Pelo^dcKO^*, see Method section for detail) (**Figure 1N**). In contrast, LPS-induced NF-κB and MAPKs activations were not affected by *Pelo* depletion (**Figure S2H** and **S2I**), and multiple Toll-like receptor (TLR) ligands mediated TNFα and IL-6 inductions were unaffected by *Pelo* deletion (**Figure S2J** and **S2K**). Thus, PELO is required for the function of NOD2 selectively.

To demonstrate the requirement of PELO in NLRP3 and NLRC4 inflammasome activation unambiguously, we reconstituted *Pelo^mKO^* BMDMs by ectopic expression of PELO. Re-expression of PELO in *Pelo^mKO^* BMDMs restored the activation of caspase-1, pyroptosis and production of IL-1β by NLRP3 and NLRC4 inflammasomes (**Figure S3A-S3C**). Of note, deletion of *PELO* also resulted in the impaired activation of caspase-1, pyroptosis and production of IL-1β in human monocyte line THP-1 cells primed with LPS and stimulated with nigericin (**Figure S3D-S3F**). Re-expression of PELO in *PELO* KO THP-1 cells restored the activation of caspase-1 and cleavage of IL-1β (**Figure S3G**).

The data aforementioned demonstrate that PELO controls the function of cytosolic NLRs. We then used NLRP3 and NLRC4 inflammasomes to further investigate the mechanism by which PELO regulates NLRs activation.

### PELO is essential for NLRP3 and NLRC4 inflammasome activation *in vivo*

As PELO participates in host defenses (**Figure 1D** and **1E**), we at first determined whether NLRP3 inflammasome activation is one of PELO regulated host responses *in vivo*. NLRP3-mediated induction of IL-1β expression in mice after intraperitoneal injection of LPS or monosodium urate crystals (MSU) and recruitment of neutrophils to the peritoneal cavity of mice after intraperitoneal injection of MSU were used as model systems (Mariathasan et al., 2006; Martinon et al., 2006; Sutterwala et al., 2006). *Pelo* deletion in the myeloid compartment of *Pelo^fl/fl^LysM^cre^* mice significantly reduced LPS-induced serum IL-1β increase in comparison with that in *Pelo^fl/fl^* mice (**Figure 2A**), whereas the induction of TNFα and IL-6 by LPS was comparable across the two groups (**Figure 2B** and **2C**). MSU-induced increase of IL-1β in the peritoneal lavage fluid (**Figure 2D**) as well as recruitment of total cells (**Figure 2E**) and neutrophils (**Figure 2F**) to the peritoneal cavity of *Pelo^fl/fl^LysM^cre^* mice were less than those of *Pelo^fl/fl^* mice.

**Figure 2.**
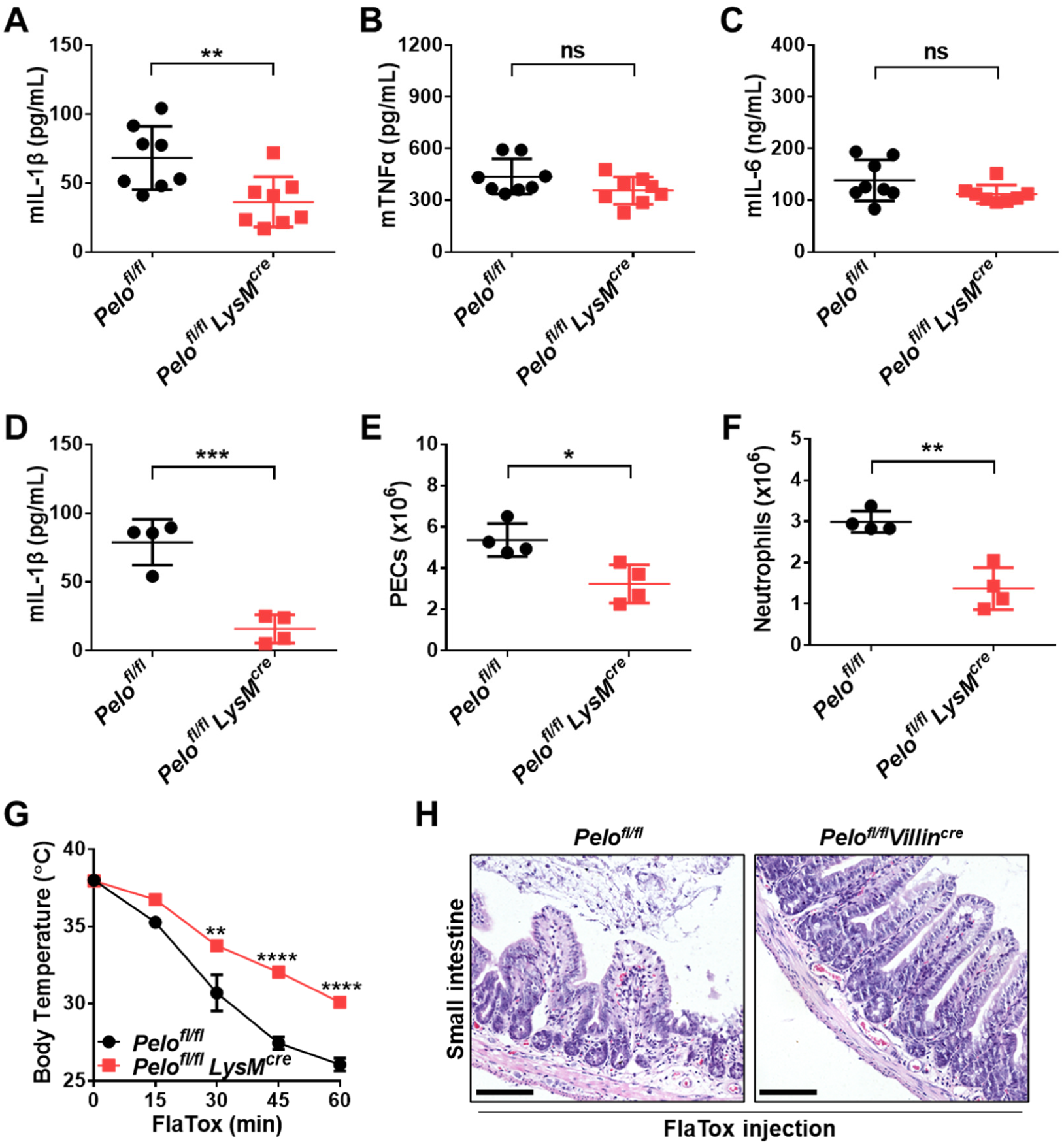
PELO is essential for NLRP3 and NLRC4 inflammasome activation *in vivo*. (**A-C**) Mouse serum cytokines IL-1β (**A**), IL-6 (**B**) and TNFα (**C**) were analyzed after an intraperitoneal injection of 10 mg/kg LPS for 3 hours. n = 8 mice per genotype. (**D-F**) Quantification of IL-1β (**D**), peritoneal exudate cells (PECs) (**E**) and neutrophils (Ly6G^+^F4/80^-^) **(F)** in the peritoneal lavage fluid of *Pelo^fl/fl^* and *Pelo^fl/fl^LysM^cre^* mice 3 hours after intraperitoneal injection of 1 mg MSU. n = 4 mice per genotype. (**G**) *Pelo^fl/fl^* and *Pelo^fl/fl^LysM^cre^* mice were intravenously injected with 0.2 μg/g PA plus 0.2 μg/g LFn-Fla (FlaTox) and monitored for body temperature. n = 4 mice per genotype. (**H**) H&E staining of small intestinal tissue from mice treated as in (**G**), 30 min after injection. Scale bars represent 100 μm. *P* values are determined by two-sided unpaired Student’s *t*-tests. ns, not significant; \**P* < 0.05; \*\**P* < 0.01; \*\*\**P* < 0.001; \*\*\*\**P* < 0.0001. Data are represented as mean ± SD (**A**-**G**). All results are representative of two independent experiments.

We also assessed the role of PELO in the regulation of NLRC4 inflammasome *in vivo*. It is known that intravenous injection of FlaTox induces an NLRC4-dependent hypothermic response, and macrophages play a key role in this response (von Moltke et al., 2012). We found that the body temperature drop was significantly attenuated in *Pelo^fl/fl^LysM^cre^* mice compared with that in *Pelo^fl/fl^* mice (**Figure 2G**). Since previous studies reported that activation of NLRC4 inflammasome in intestinal epithelial cells (IECs) led to small intestine damages (Rauch et al., 2017; Zhang et al., 2021), we analyzed the small intestines in *Pelo^fl/fl^Villin^cre^* mice and observed that deletion of *Pelo* protected IECs from FlaTox-induced sloughing and villus blunting in the small intestines (**Figure 2H**). Thus, PELO is essential for NLRP3 and NLRC4 inflammasome activation *in vivo*.

### PELO promotes inflammasome activation independent of ribosome rescue

Because PELO is an essential player in ribosome rescue, we next investigated whether PELO’s function in inflammasomes links to ribosome rescue. PELO forms a complex with HBS1L to rescue the stalled ribosomes (Carr-Schmid et al., 2002; Doma and Parker, 2006; Guydosh and Green, 2014; Pisareva *et al*., 2011; Shoemaker *et al*., 2010; Tsuboi *et al*., 2012). Previous studies showed that PELO and HBS1L stabilized each other in mouse tissues and primary mouse embryonic fibroblasts (MEFs) (O’Connell et al., 2019; Terrey *et al*., 2021), and we found *Pelo* deficiency also led to a decrease of HBS1L protein in BMDMs and RAW-ASC cells (RAW264.7 macrophage cell line ectopically expressing ASC (He *et al*., 2015))(**Figure S4A**). To determine whether the decrease of HBS1L plays a role in inflammasome activation, we restored the expression of HBS1L in *Pelo^mKO^* BMDM cells. As shown in **Figure S4B**, re-expression of PELO was able to fully restore both the expression of HBS1L and the cellular response to NLRP3 and NLRC4 inflammasome activators, whereas re-expression of HBS1L could not. To further determine the requirement of HBS1L in NLRP3 or NLRC4 activation, we generated *Hbs1l^-/-^* chimeric mice (**see Method**). We obtained three chimeric mice with no detectable level of HBS1L protein in their bone marrow (**Figure S4C)**. The deficiency of *Hbs1l* in BMDMs did not alter the activation of caspase-1, pyroptosis and maturation of IL-1β by NLRP3 or NLRC4 inflammasome activation (**Figure 3A-3C**). The activation of NLRP3 inflammasome was not affected in *Hbs1l* KO RAW-ASC cells and this cell line with additional deletion of *Gtpbp2*, a recently identified binding partner of PELO that functions in the ribosome rescue process (Ishimura et al., 2014) (**Figure 3D-3F**). Thus, there is no dependence between PELO-mediated regulation of NLRs and ribosome rescue.

**Figure 3.**
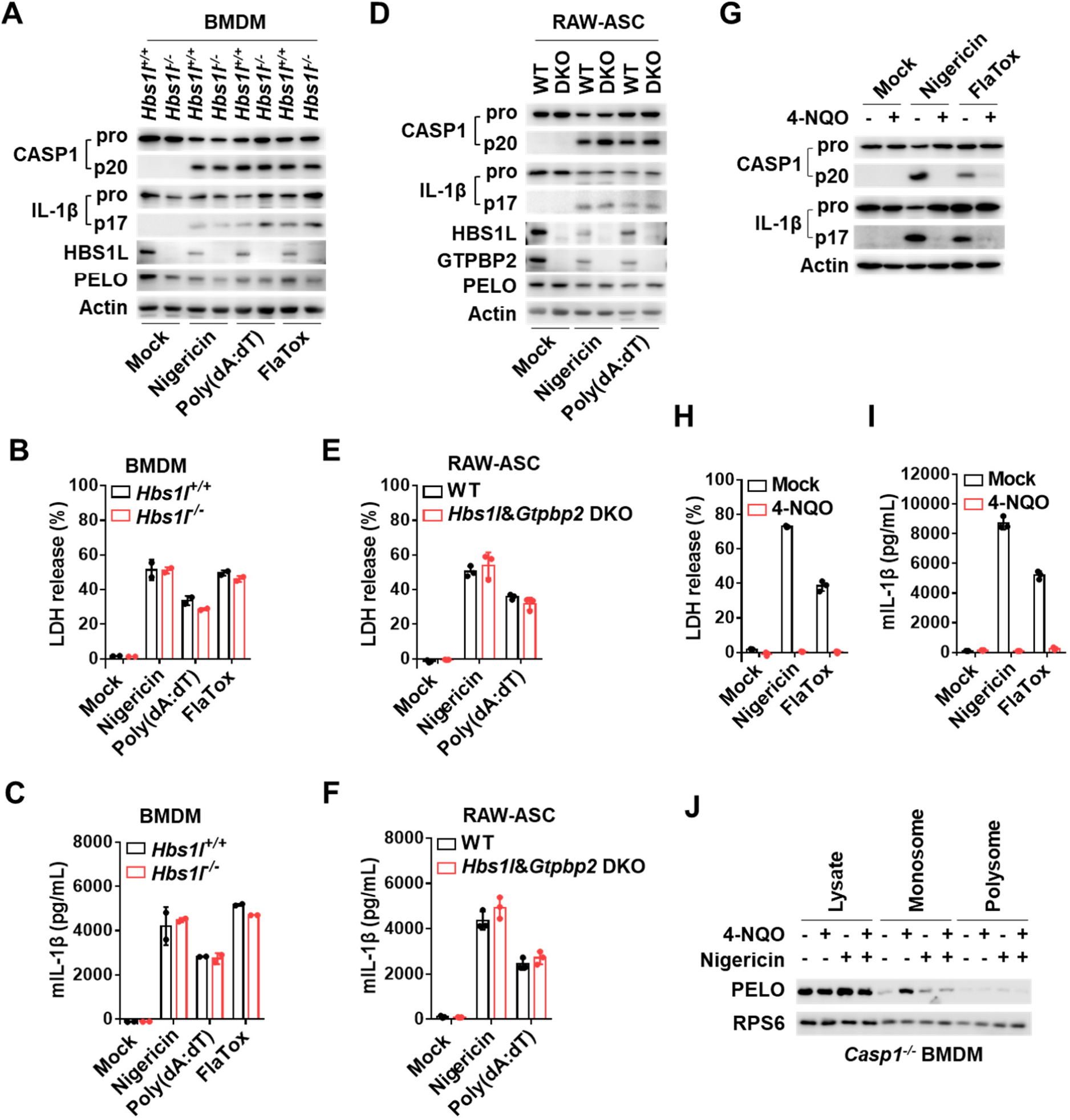
PELO promotes inflammasome activation independent of ribosome rescue, but these two processes could compete for PELO protein. (**A**) Immunoblot analysis of the processed caspase-1 (CASP1) and IL-1β in the pooled cell extracts and supernatants from *Hbs1l^+/+^* and *Hbs1l^-/-^* BMDMs that were primed with LPS and then treated with the indicated stimuli. (**B** and **C**) BMDMs were treated as in (**A**); LDH (**B**) and IL-1β (**C**) in culture supernatants were analyzed. Data are represented as mean ± SD of triplicate wells. (**D**) Immunoblot analysis of the processed caspase-1 (CASP1) and IL-1β in the pooled cell extracts and supernatants from WT and *Hbs1l* plus *Gtpbp2* double knockout (DKO) RAW-ASC cells that were primed with LPS and then treated with the indicated stimuli. (**E** and **F)** RAW-ASC cells were treated as in (**D**). LDH (**E**) and IL-1β (**F**) in culture supernatants were analyzed. Data are represented as mean ± SD of triplicate wells. (**G-I**) LPS-primed BMDMs were left untreated or pre-treated with 4-NQO (20 μM) for 30 min and then treated with the indicated stimuli. The processed caspase-1 (CASP1) and IL-1β in the pooled cell extracts and supernatants were analyzed by immunoblotting (**G**); LDH (**H**) and IL-1β (**I**) in culture supernatants were measured. Data are represented as mean ± SD of triplicate wells. (**J**) LPS-primed *Casp1^-/-^*BMDMs were left untreated or pre-treated with nigericin for 30 min and then treated with 4-NQO. The polysome and monosome fractions were analyzed by immunoblotting as indicated. All results are representative of three independent experiments. See also **Figure S4** and **S5**.

### Inflammasome activation and ribosome rescue could compete for PELO

UV-mimic 4-nitroquinoline 1-oxide (4-NQO), which generates ROS and oxidizes guanosine nucleobase (8-oxoguanosine, 8-oxo-G), can stall ribosomes by altering codon-anticodon interactions and thus activates the PELO-dependent ribosome rescue process (Simms et al., 2014; Yan et al., 2019). As expected, ribosome profiling revealed that PELO was predominantly found with the light fractions, and 4-NQO treatment moved a significant amount of PELO to monosome fractions (**Figure S4D** and **S4E**). To stall ribosomes in LPS-primed BMDMs, we treated them with 4-NQO, and then stimulated them with nigericin or FlaTox. Interestingly, the cleavage of caspase-1 and pyroptosis, as well as cleavage of pro-IL-1β and release of IL-1β were strongly inhibited by 4-NQO (**Figure 3G-3I**). This inhibition was primarily on the inflammasome assembly since the expression levels of pro-IL-1β, NLRP3, NLRC4, ASC and CASP1 were not affected (**Figure S4F**). However, 4-NQO also has some suppressive effects on LPS-induced TNFα and IL-6 expression, indicating 4-NQO might has an effect other than ribosome stalling (**Figure S4G** and **S4H**). We also tested if 4-NQO exerted an impact on the formation of G3BP1 and DDX3X containing stress granules that can inhibit NLRP3 inflammasome activation (Samir et al., 2019) and found no effect (**Figure S4I** and **S4J**). When 4-NQO was added at different time points before or after nigericin or FlaTox stimulation of LPS-primed BMDMs, later 4-NQO treatment had less or no effect on the activation of NLRP3 and NLRC4 inflammasomes (**Figure S4K** and **S4L**). To induce the assembly of NLRP3 inflammasome before 4-NQO treatment, we used *Casp1^-/-^*BMDMs to avoid pyroptosis. We found that the already formed inflammasomes inhibited 4-NQO induced movement of PELO to the monosome fractions (**Figure 3J**). Collectively, our data suggest that ribosome stalling and NLRP3 inflammasome activation compete for the use of PELO. This notion is supported by the fact that PELO is a low-abundance protein, whose copy number per cell is less than 88% of the other cellular proteins in BMDMs (**Table S2**).

We also analyzed the effect of PELO on NLRP1 inflammasome, whose activation can be triggered by the degradation of its PYD, NACHT and LRR domain containing N-terminal fragment, and the degradation is often initiated by microbial effector proteases (Chui et al., 2019; Robinson et al., 2020; Sandstrom et al., 2019). NLRP1B variants from BALB/c and 129 mouse strains can be activated by the lethal toxin of *Bacillus anthracis* that is comprised of lethal factor (LF) protease and protective antigen (PA) (Boyden and Dietrich, 2006). The role of PA is to deliver LF into the host cells. RAW264.7 cell was a macrophage cell line derived from BALB/c mouse strain, and *Pelo* deletion had no effect on the activation of NLRP1B inflammasome induced by anthrax lethal toxin in RAW-ASC cells (**Figure S5A-S5F**). This result is expected since PELO interacts with NLRP1B *via* the N-terminal portion of NLRP1B (**Figure S5G**) and the removal and subsequent degradation of the N-terminal fragment left no chance for PELO to promote NLRP1B activation. Consistently, the PELO-independent activation of NLRP1B inflammasome by lethal toxin was not affected by 4-NQO treatment (**Figure S5H-S5J**).

### PELO is required for the activation of NLR but not non-NLR inflammasomes

Formation of large intracellular ASC aggregates (called ASC specks) in cytosol is a hallmark of complete activation of inflammasomes (Fernandes-Alnemri et al., 2007). As expected, stimulation of BMDMs with NLRP3 activators nigericin or R837, NLRC4 activators FlaTox or *Salmonella,* or AIM2 activator poly(dA:dT), induced the formation of ASC specks in cytosol (**Figure 4A** and **4B**). PELO deficiency dramatically reduced the formation of ASC specks induced by NLRP3 and NLRC4 inflammasomes, whereas it did not affect the formation of ASC specks induced by poly(dA:dT) (**Figure 4A** and **4B**). Consistently, analysis of ASC oligomers in detergent-insoluble fraction revealed that NLRP3- and NLRC4-but not AIM2-mediated ASC oligomerization was impaired in *Pelo^mKO^* BMDMs (**Figure 4C**). These data together with the data in **Figure 1C**, **1F**, **1G**, **1H** and **1L** showed that PELO is specifically required for the activation of those NLR inflammasomes.

**Figure 4.**
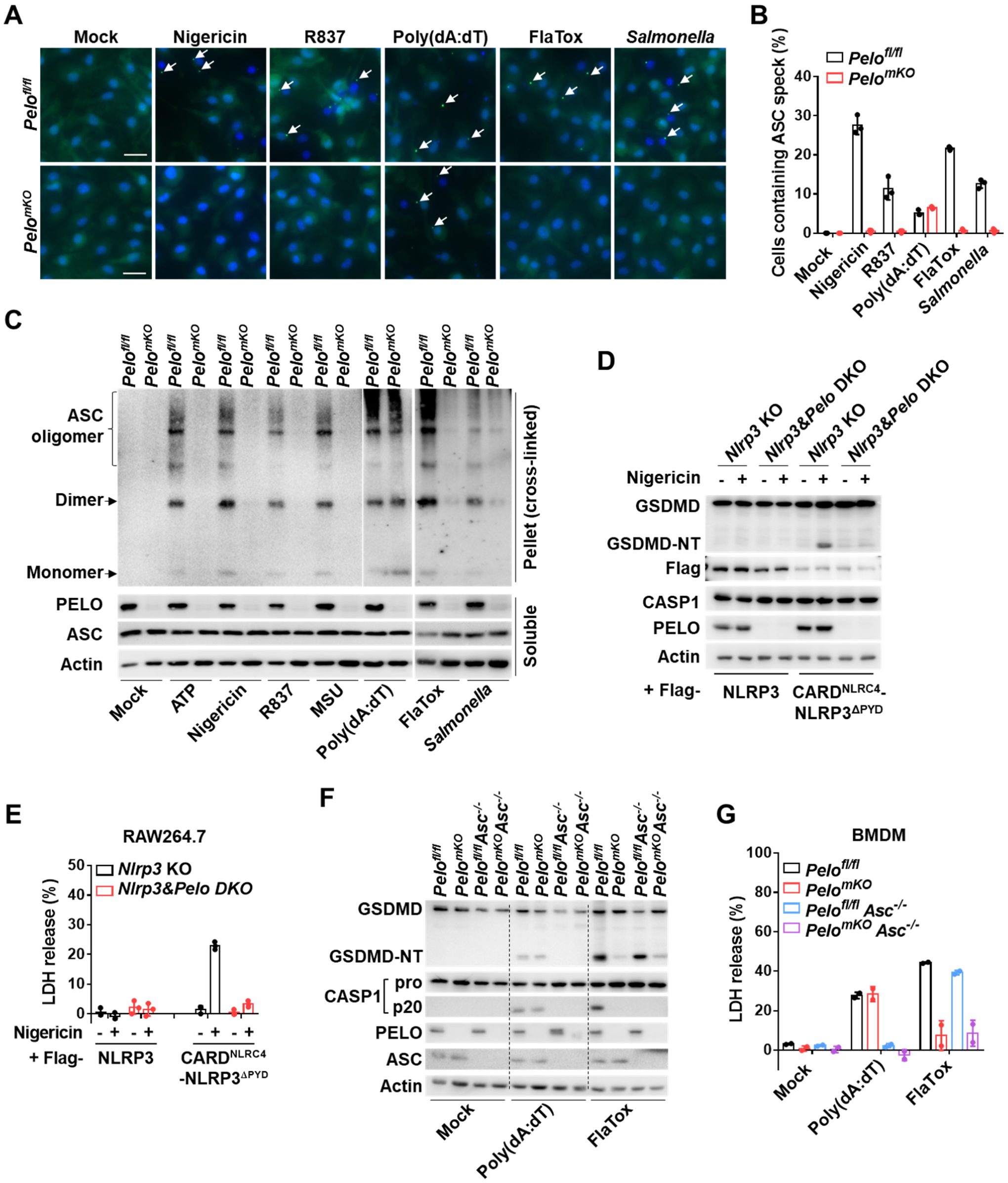
PELO functions at upstream of ASC speck formation and is specifically required for NLR inflammasomes. (**A** and **B**) Representative immunofluorescence images and quantification of ASC specks (arrows) in *Pelo^fl/fl^* and *Pelo^mKO^* BMDMs that were primed with LPS and then treated with the indicated stimuli. Scale bars represent 20 μm. Data are represented as mean ± SD (n = 3 biologically independent fields of cells). (**C**) Immunoblot analysis of ASC oligomerization induced by indicated stimuli as in (**A**). The samples were cross-linked (see the method section). (**D** and **E**) *Nlrp3* knockout (KO) and *Nlrp3* plus *Pelo* double knockout (DKO) RAW264.7 cells with ectopic expression of Flag-tagged WT NLRP3 or CARD^NLRC4^-NLRP3^ΔPYD^ chimera were stimulated with nigericin for 1 hour. The cleavage of GSDMD in the pooled cell extracts and supernatants (**D**) and LDH in the supernatants (**E**) were analyzed. Data are represented as mean ± SD of triplicate wells. (**F** and **G**) BMDMs from mice of the indicated genotypes were stimulated as indicated. The cleavage of GSDMD in the pooled cell extracts and supernatants (**F**) and LDH release from cells (**G**) were analyzed. n = 2 mice per genotype. Data are represented as mean ± SD. All results are representative of at least two independent experiments.

ASC is a required adaptor for NLRP3 but dispensable for NLRC4 to activate caspase-1 due to that the CARD domain in NLRC4 allows its direct interaction with caspase-1 (Broz et al., 2010; Mariathasan et al., 2004). This implies that ASC might not be required for PELO to function in inflammasome activation. To explore whether ASC is required for PELO to regulate NLRP3 inflammasome, we adapted a reported strategy that replacement of the pyrin domain (PYD) of NLRP3 with the CARD domain of NLRC4 (CARD^NLRC4^-NLRP3^ΔPYD^ chimera) can result in NLRP3 inflammasome activation in the absence of ASC (Hafner-Bratkovic et al., 2018). We expressed Flag-CARD^NLRC4^-NLRP3^ΔPYD^ and Flag-NLRP3 in *Nlrp3* KO RAW264.7 cells (which do not express ASC) and confirmed that CARD^NLRC4^-NLRP3^ΔPYD^ but not NLRP3 can mediate nigericin-induced GSDMD cleavage and cell death (**Figure 4D** and **4E**). Deletion of *Pelo* greatly reduced nigericin-induced GSDMD cleavage and cell death in CARD^NLRC4^-NLRP3^ΔPYD^ chimera reconstituted RAW264.7 cells (**Figure 4D** and **4E**), indicating that PELO functions in a step other than ASC speck formation in NLRP3 activation.

In the case of NLRC4 inflammasome activation, deletion of *Asc* in BMDMs eliminated the ASC-dependent cleavage of caspase-1 but GSDMD was still cleaved during NLRC4 inflammasome activation and cell death still occurred (**Figure 4F** and **4G**), which is expected as that although caspase-1 did not auto-process in the absence of ASC, the full-length caspase-1 recruited by NLRC4 still can process a certain amount of GSDMD and cause cell death (Broz *et al*., 2010). Since deletion of *Pelo* significantly inhibited NLRC4-induced GSDMD cleavage and cell death regardless of the presence of ASC (**Figure 4F** and **4G**), PELO has no direct regulatory role on ASC speck formation.

### PELO promotes the oligomeric assembly of NLRP3 and NLRC4 inflammasomes

To this point, our results collectively pointed out that PELO should function at the initial step of inflammasome assembly. We then confirmed that PELO-NLRP3 interaction associates with the initiation of NLRP3 inflammasome assembly, as the interaction between endogenous PELO and NLRP3 in LPS-primed BMDMs was induced after nigericin treatment (**Figure 5A**). Due to a lack of good NLRC4 antibody for immunoprecipitation, we reconstituted *Nlrc4^-/-^* BMDMs with Flag-tagged NLRC4. The interaction between endogenous PELO and Flag-NLRC4 was substantially enhanced by stimulation with FlaTox (**Figure 5B**). We also detected NLRP3-PELO interaction and NLRC4-PELO interaction in Flag-PELO-reconstituted BMDMs by anti-Flag immunoprecipitation (**Figure 5C** and **5D**).

**Figure 5.**
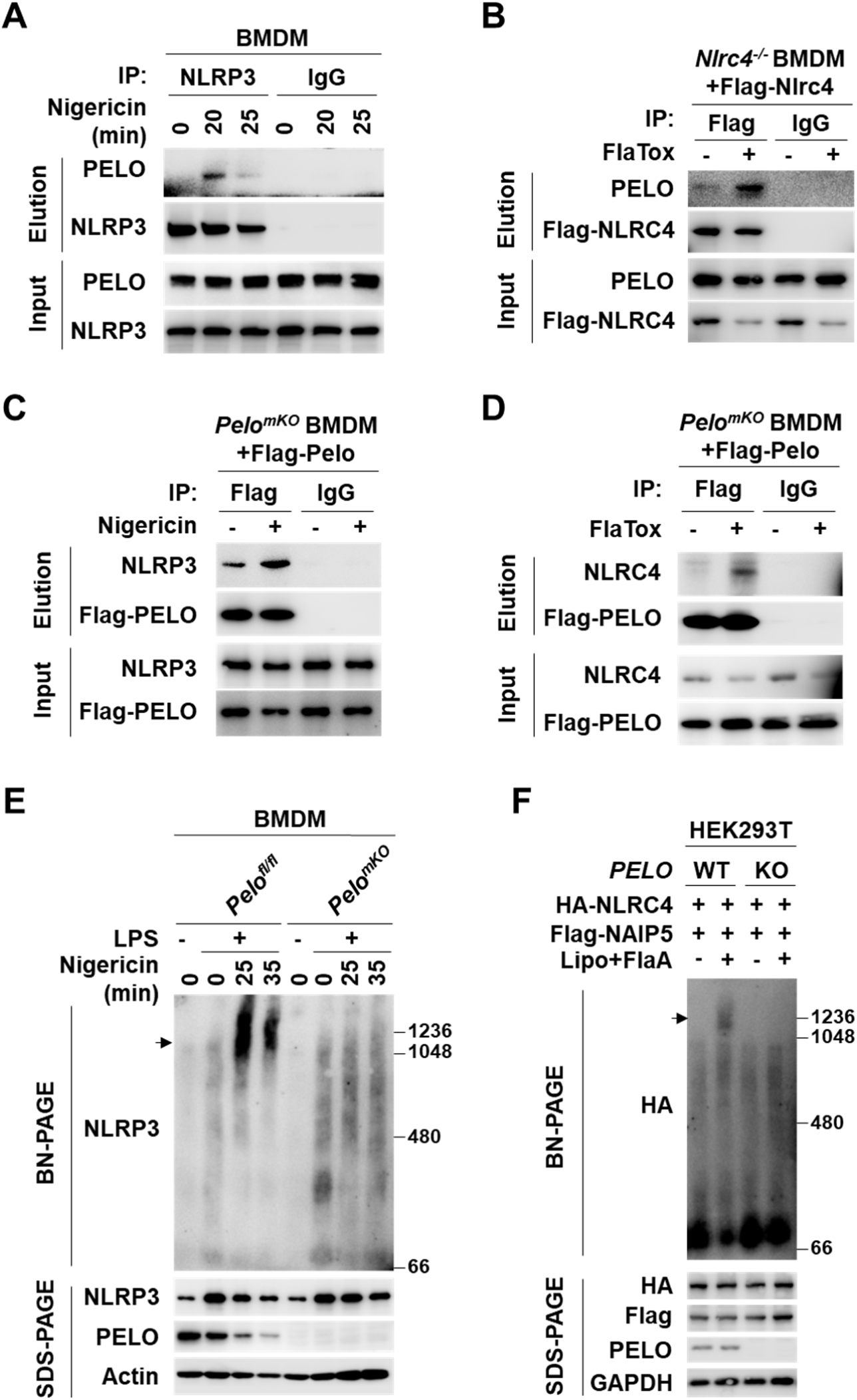
PELO promotes the oligomeric assembly of NLRP3 and NLRC4 inflammasomes in cells. (**A**) LPS-primed BMDMs were stimulated with nigericin as indicated. The cell lysates were immunoprecipitated with anti-NLRP3 or IgG antibodies, followed by immunoblot analysis. (**B**) *Nlrc4^-/-^* BMDMs reconstituted with Flag-NLRC4 were treated with FlaTox for 30 min and the cell lysates were immunoprecipitated with anti-Flag or IgG antibodies, followed by immunoblot analysis. (**C** and **D**) *Pelo^mKO^* BMDMs reconstituted with Flag-Pelo were treated with LPS (4 hours) plus nigericin (30 min) (**C**) or FlaTox (30 min) (**D**) as indicated and the cell lysates were immunoprecipitated with anti-Flag or IgG antibodies, followed by immunoblot analysis as indicated. (**E**) *Pelo^fl/fl^* and *Pelo^mKO^* BMDMs were left untreated or primed with LPS for 4 hours and then stimulated with nigericin for indicated time periods. The cell lysates were resolved by blue native PAGE (BN-PAGE) for detection of NLRP3 in high-molecular-weight complexes (arrow) or by SDS-PAGE for detection of total NLRP3 (bottom), followed by immunoblot analysis. (**F**) Wild-type (WT) and *PELO* KO HEK293T cells were transfected with expression vectors of Flag-NAIP5 together with HA-NLRC4. 24 hours later, the cells were transfected with or without flagellin (FlaA) protein by Lipo2000. 6 hours later, the cell lysates were analyzed as in (**E**). Note: HEK293T cell has no endogenous expression of NLRP3, NLRC4 or NAIP5. All results are representative of at least two independent experiments. See also **Figure S6**.

To determine the involvement of PELO in the oligomerization of NLRP3 or NLRC4, we analyzed inflammasomes in *Pelo^fl/fl^* and *Pelo^mKO^* BMDMs. The digitonin-solubilized cell lysates with the insoluble ASC specks having been removed by ultracentrifugation were resolved by blue native PAGE (BN-PAGE). A large oligomeric complex (>1,000 kDa) containing NLRP3 was induced in *Pelo^fl/fl^* BMDMs after stimulation with nigericin, but the formation of this complex was greatly diminished in *Pelo^mKO^* BMDM cells (**Figure 5E**). To visualize NLRC4 inflammasome assembly, NAIP5 and NLRC4 were expressed in HEK293T cells, and as reported (Kofoed and Vance, 2011), a shift of NLRC4 from a monomer (∼120 kDa) to an oligomeric complex (∼1,200 kDa) can be observed when flagellin was co-expressed (**Figure S6A**) or when flagellin was delivered into cells by Lipo2000 or PA (**Figure 5F** and **S6B**). Deletion of *PELO* eliminated the oligomerization of NLRC4 in this assay (**Figure 5F**, **S6A** and **S6B**). Thus, PELO participates in NLRP3- or NLRC4-oligomerization during inflammasome activation.

As NEK7 and DDX3X have been shown to mediate NLRP3 inflammasome assembly and activation (He et al., 2016; Samir *et al*., 2019; Schmid-Burgk et al., 2016; Shi et al., 2016), we further addressed how these identified NLRP3 partners orchestrate NLRP3 inflammasome activation together with PELO. We compared the NLRP3 complex between *Pelo^fl/fl^* and *Pelo^mKO^* BMDMs by immunoprecipitation, and detected inducible recruitments of NEK7, DDX3X, as well as PELO to the NLRP3 complex in *Pelo^fl/fl^* BMDMs (**Figure S6C**). Notably, we found that deletion of *Pelo* abolished the recruitment of NEK7 and DDX3X to NLRP3 (**Figure S6C**). Quenching PELO by 4-NQO-induced ribosome rescue severely impaired the formation of nigericin-induced NLRP3 complex (**Figure S6D**). As expected, BMDMs derived from *Nek7^fl/fl^LysM^cre^* mice (*Nek7^mKO^* BMDMs) showed impairment of NLRP3 inflammasome-mediated pyroptosis similar to that of *Pelo^mKO^* BMDMs (**Figure S6E**). Moreover, the absence of NEK7 also blocked the interaction of PELO and DDX3X with NLRP3 (**Figure S6F**), indicating that both PELO and NEK7 are necessary and non-redundant for functional NLRP3 complex assembly.

### PELO promotes NLRC4 inflammasome assembly *via* activating the ATPase of NLRC4

Successful reconstitution of the inflammasome by purified flagellin, NAIP5, and NLRC4 *in vitro* has been reported (Halff et al., 2012), which provides a system to address the mechanism by which PELO promotes the oligomeric assembly of NLR proteins. We expressed HA-NLRC4 and Flag-NAIP5 respectively or together in *PELO* KO HEK293T cells, and observed that these two proteins were in their monomer form in the cell lysates but NLRC4 became aggregates with various sizes after purifying NLRC4 from cell lysates. The aggregation of NLRC4 was observed in the previous study when the concentration of NLRC4 protein was high but NAIP5 had no tendency to self-aggregate (Halff *et al*., 2012). We thus directly used cell extracts from HA-NLRC4 and Flag-NAIP5 co-expressing *PELO* KO HEK293T cells as the source of NLRC4 monomer protein and Flag-NAIP5 for the *in vitro* experiments. HA-NLRC4 and Flag-NAIP5 were mixed and incubated with or without recombinant flagellin plus ATP *in vitro*, but the HA-NLRC4 oligomerization was not detected by BN-PAGE analysis of the reaction mixtures containing NLRC4 and NAIP5 with the supplement of flagellin+ATP (**Figure 6A**, Lane 4). Previously reported *in vitro* assembly of NLRC4 inflammasome by the three proteins: flagellin, NAIP5 and NLRC4, was known to be low efficient (Halff *et al*., 2012), and here was not detectable in our system. Nonetheless, we were going to address the role of PELO and therefore included PELO in the reaction. Notably, the addition of PELO protein resulted in efficient formation of a high molecular weight complex with a similar apparent molecular weight to the NAIP5-NLRC4 complexes from flagellin stimulated cells (**Figure 6A**, Lane 6). Thus, flagellin, ATP, and PELO are all required for efficient NAIP5-NLRC4 oligomerization.

**Figure 6.**
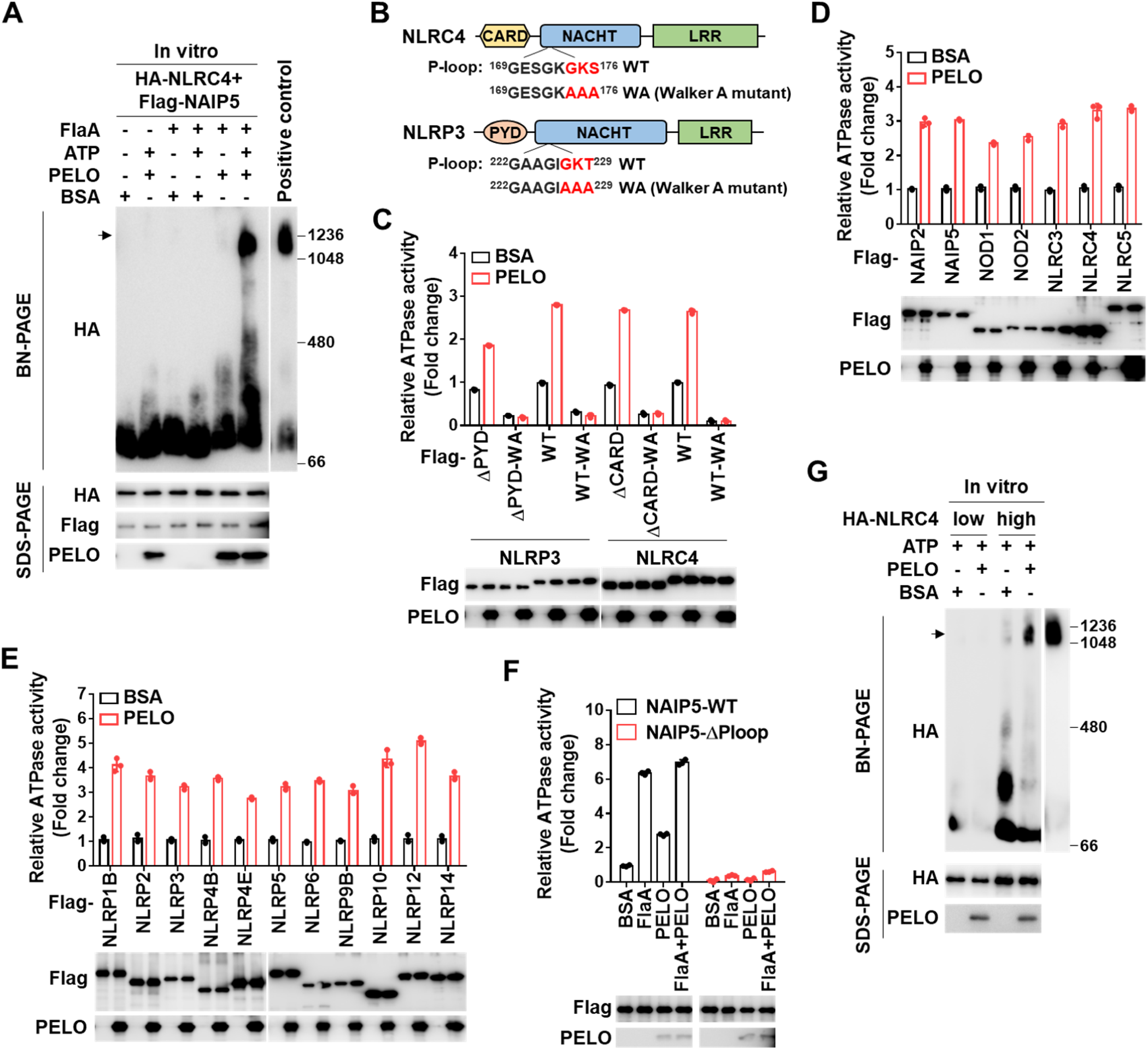
PELO promotes NLRC4 inflammasome assembly *via* activating the ATPase of NLRC4. (**A**) HA-NLRC4 and Flag-NAIP5 containing *PELO* KO HEK293T cell lysates were incubated with purified recombinant flagellin (FlaA), ATP and recombinant His-PELO in different combinations as indicated. BSA was used as the control of PELO. After 1 hour of incubation at 37℃, the mixtures were analyzed by BN-PAGE. (**B**) Schematics of the structures of NLRC4 and NLRP3 and the sequence of P-loop region and Walker A mutation. (**C**) ATPase activity was measured by a bioluminescent assay using purified Flag-WT or mutant NLRP3 and NLRC4 in the presence of BSA or recombinant PELO protein. (**D** and **E)** ATPase activities of purified Flag-NLR family proteins in the presence of BSA or recombinant His-PELO protein. Folds of change were shown. (**F**) ATPase activity of purified Flag-WT or mutant NAIP5 in the presence of BSA, FlaA or PELO protein. Folds of change were shown. Data are represented as mean ± SD of triplicates (**C-F**). (**G**) Cell lysates containing low or high concentrations of HA-NLRC4 were incubated with ATP and with or without PELO as indicated and analyzed by BN-PAGE. All results are representative of at least two independent experiments. See also **Figure S7**.

The current model for NLRC4 activation is that flagellin binds to NAIP5, which by itself cannot oligomerize but induces the recruitment and progressive incorporation of NLRC4 to form oligomeric inflammasome, and the complex assembly requires hydrolysis of ATP by activation of ATPase. Consistent with published studies (Kofoed and Vance, 2011), the Walker A (WA) mutant (ATP binding defect) of NLRC4 (**Figure 6B**) abolished the formation of NLRC4 oligomer both in flagellin-stimulated cells (**Figure S7A**) and in reconstitution assay *in vitro* (**Figure S7B**). Since PELO had no effect on the assembly of NLRC4 inflammasome in the absence of ATP (**Figure 6A**) and PELO interacts with NACHT domain, PELO’s function might link to ATPase activity of NLRC4 and/or NAIP5.

It is known that PELO increases the GTPase activity of its partner HBS1L in the ribosome rescue pathway (Chen et al., 2010; Graille et al., 2008; Pisareva *et al*., 2011; Shoemaker *et al*., 2010). And the sequence conservation has placed NLRs as members of the STAND AAA+ ATPase superfamily (Lelpe *et al*., 2004). The central NACHT domain in all NLRs bears conserved ATP-binding and hydrolysis motifs (Koonin and Aravind, 2000; Lelpe *et al*., 2004), and activated STAND proteins are known to promote oligomeric assemblies by coupling ATP hydrolysis to drive conformational reorganization of the protein structure and surface exposure of previously concealed binding sites for multimeric complex formation (Danot *et al*., 2009). The ATPase activity of NLRP3, NLRC4 and several other NLRs had been shown to be essential for their oligomerization and activation (Sandall et al., 2020). Since PELO interacts with the NACHT domain of NLRP3 and NLRC4 (**Figure S1B** and **S1C**), we sought to test whether PELO could affect the ATPase activity of NLRP3 and NLRC4. To assess the ATPase activity of NLRP3 and NLRC4, we purified NLRP3, NLRC4 and their mutants from HEK293T cells **(Figure S7C)**. Each of the purified proteins was incubated with ATP and then the production of ADP was determined by a bioluminescent reporter assay. NLRP3 and NLRC4 but not their Walker A (WA) mutants exerted ATPase activity as previously reported (Duncan et al., 2007; Lu et al., 2005) (**Figure 6B** and **6C**). Importantly, addition of purified recombinant PELO protein into the incubation significantly increased ATPase activity of NLRP3 and NLRC4 but had no effect on their WA mutants (**Figure 6B** and **6C**). As expected, deletion of PYD or CARD domain had no influence on the ATPase activity of NLRP3 or NLRC4 (**Figure 6B** and **6C**), and the Walker A mutation in NLRP3 and NLRC4 did not affect their interactions with PELO (**Figure S7D**). Notably, the effect of PELO on the ATPase activity also applies to all the other cytosolic NLRs (**Figure 6D**, **6E**, **S7E** and **S7F**). Given the fact that the ATPase activity is required at least for certain NLRs to function (Sandall *et al*., 2020), potentiation of ATPase activity of a given NLR by PELO should be necessary for the oligomerization and activation of this NLR.

NLRC4 inflammasome assembly is known to be initiated by binding of flagellin to NAIP5. Previous studies had different conclusions on whether the ATPase activity of NAIP5 is required for NLRC4 inflammasome assembly (Halff *et al*., 2012; Kofoed and Vance, 2011). We re-visited these studies and concluded that the confusion was most likely caused by different mutants used. The K476R mutation in the Walker A motif of NAIP5 retained a low level-ATPase activity (**Figure S7G** and **S7H**), and thus only partially blocked flagellin-induced NLRC4 oligomerization (**Figure S7I**). These data led us to agree that the ATPase activity of NAIP5 is required for initiating NLRC4 inflammasome activation, and then we analyzed the role of flagellin on the ATPase activity of NAIP5. The ATPase activity of NAIP5 was stimulated by flagellin, and further including PELO in the incubation can only slightly enhance the ATPase activity (**Figure 6F**). Thus, PELO is most likely dispensable for the activation of NAIP5 by flagellin. Since flagellin cannot interact with NLRC4, we asked whether PELO can promote NLRC4 oligomerization. NLRC4 protein tended to aggregate *in vitro* unless its concentration was low (Halff *et al*., 2012). We incubated NLRC4 at different concentrations in the presence or absence of PELO, and found that PELO caused oligomerization of NLRC4 to the sizes of functional NLRC4 complex (**Figure 6G**). Thus, PELO should promote NLRC4 inflammasome assembly *via* activating the ATPase of NLRC4 but not NAIP5.

### PELO functions as a catalyzer of NLRC4 inflammasome assembly

Structural studies have revealed that the NLRC4 inflammasomes are disk-shaped particles containing 10-12 symmetric protomers, and the stoichiometry of the ligand, NAIP, and NLRC4 constituents within inflammasome complexes was estimated to be around 1:1:10 (Hu et al., 2015; Zhang et al., 2015). Indeed, about a 1:10 ratio of NAIP5 to NLRC4 was observed in the NLRC4 complex purified by flagellin pulldown in our experimental system (**Figure 7A**). Importantly, the PELO in the complex was much less than NAIP5 and its molar ratio to NLRC4 was estimated less than 1 to 20 (**Figure 7A**). We again estimated the ratio of PELO to NLRC4 by dilution of NLRC4 complexes purified by NAIP5 pulldown, and confirmed it was around 1: 20 (**Figure 7B**). In an independent approach, the NAIP5, NLRC4 and PELO co-expressing HEK293T cells were stimulated with flagellin and the cell lysates were fractionated by sucrose gradient ultracentrifugation. We observed a clear shift of NLRC4 from light fraction to heavy fraction after flagellin stimulation (**Figure 7C**). The ratio among flagellin, NAIP5, NLRC4, and PELO in the shifted fractions varied and the proportion of PELO was much lower in the peak fraction 6 (**Figure 7C**), suggesting a dynamic association of PELO with NLRC4 complexes and possible release of PELO from completely assembled inflammasomes.

**Figure 7.**
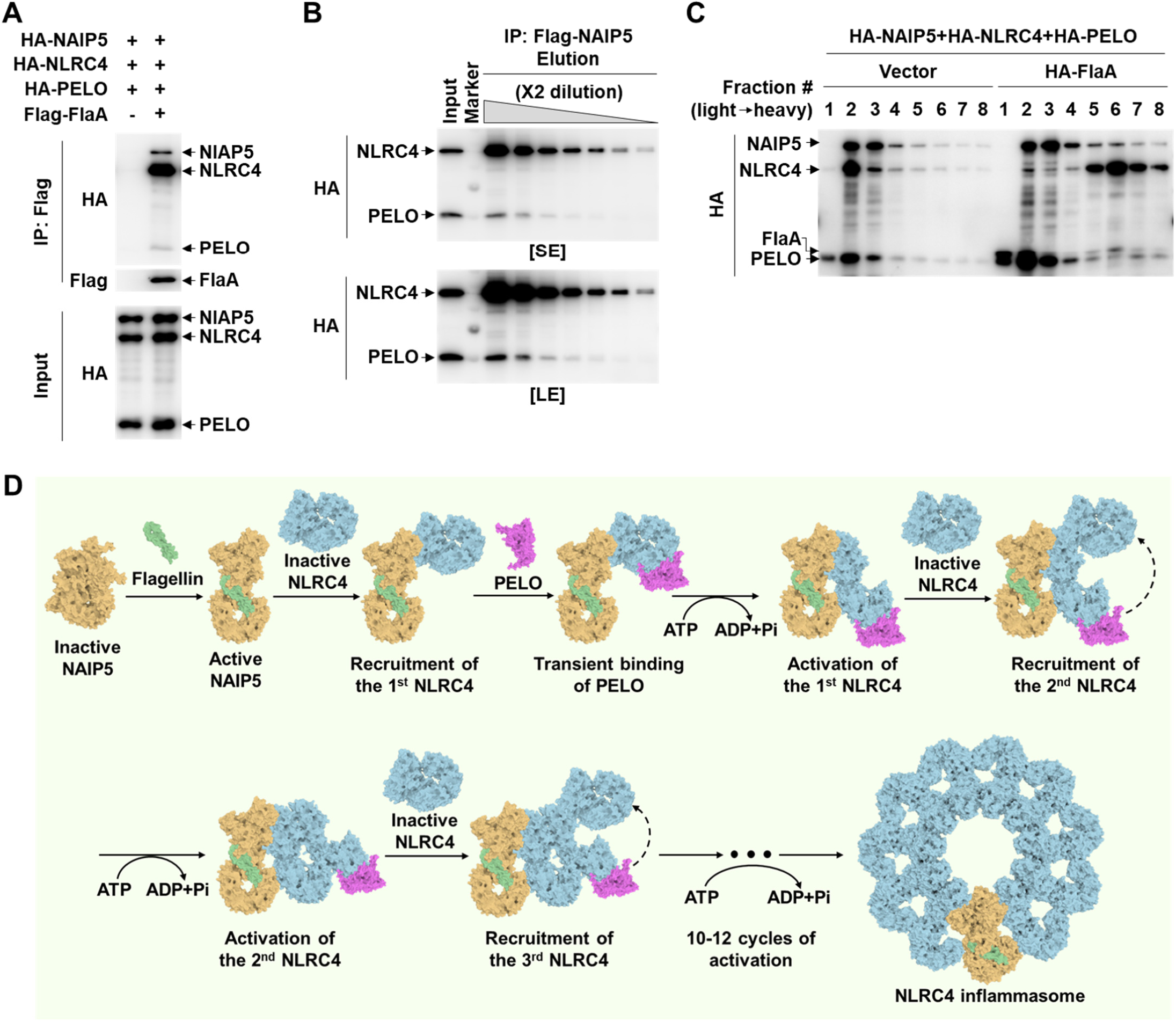
PELO acts as a catalyzer of oligomeric assembly of NLRC4. (**A**) HA-tagged NAIP5, NLRC4 and PELO were co-expressed with or without Flag-tagged flagellin (FlaA) in HEK293T cells. The cell lysates were immunoprecipitated with anti-Flag antibodies, and then the cell lysates and immunoprecipitates were analyzed by immunoblotting as indicated. (**B**) Flag-tagged NAIP5, HA-tagged NLRC4 and PELO were co-expressed with FlaA in HEK293T cells. The cell lysates were immunoprecipitated with anti-Flag antibodies, and then the cell lysates and immunoprecipitates (with 2-fold series dilution) were analyzed by immunoblotting as indicated. (**C**) HA-tagged NAIP5, NLRC4 and PELO were co-expressed with or without HA-tagged FlaA in HEK293T cells. The cell lysates were further fractionated by sucrose gradient ultracentrifugation followed by immunoblotting of each fraction as indicated. (**D**) A schematic diagram for the flagellin-induced assembly of the dish-like structure of a flagellin-NAIP5-NLRC4 complex. See text for detail. All results are representative of at least two independent experiments.

The overall around 1:20 molar ratio of PELO to NLRC4 and the non-uniform distribution of PELO in NLRC4 complexes indicate PELO is not a structural constituent of the 10 to 12-mer NLRC4 disk-shaped inflammasome particles. Hydrolysis of ATP by PELO-dependent ATPase in NLRC4 is most likely to drive the NLRC4 conformational changes, which are required for the newly incorporated NLRC4 to adapt to the right structure and for the progressive recruitment of additional NLRC4 to complete the 10 to 12-mer dish-like complex. Thus, PELO can be best characterized as a catalyzer during the assembly of NLRC4 inflammasomes. With all the information in mind, we proposed a working model for NLRC4 inflammasome assembly (**Figure 7D**). After flagellin binds to NAIP5, the activated NAIP5 initiates the recruitment of NLRC4. A PELO may transiently associate with this recruited NLRC4 and activate its ATPase, driving its conformational change to fit in the complex with right structure and then to recruit another NLRC4 to the end of the oligomerization chain of the assembling complex. The PELO might simultaneously/then move to and activate the newly recruited NLRC4. Once the oligomerization ends with the seal of the disk, PELO could release from the completely assembled NAIP5-NLRC4 10 to 12-mer disk complexes. This catalyzer model differs from the most commonly used auto-inhibition (lock) and ligand binding-mediated release of inhibition (open) model in that an additional catalyzer is required for the receptor activation besides ligand. Given the fact that PELO can bind to and potentiate the ATPase activity of all the cytosolic NLR proteins (**Figure 1C**, **6D** and **6E**), PELO could act as a catalyzer to promote the ATPase activity-driven oligomeric assembly and activation of all cytosolic NLR proteins.

## DISCUSSION

The NLR family of proteins are well known as important players in innate immune responses (Babamale and Chen, 2021; Chen *et al*., 2009; Geddes *et al*., 2009; Meunier and Broz, 2017). PELO is the only protein found to date that could participate in the activation of all cytosolic NLRs. PELO might act as a catalyzer to potentiate the NLRs ATPase activity through direct interaction with their NACHT and/or LRR domain(s). Analogous to other members of the STAND ATPase family, the hydrolysis of ATP by its activated ATPase of a given NLR protein shall remove the typic auto-inhibition structure by reorganizing its surface exposure of concealed binding sites, allowing the progressive recruiting of NLRs to form a multimeric complex. PELO appears to be selectively involved in promoting the oligomerization of NLR family members, as its deficiency had no effect on staurosporine-induced intrinsic (Apaf-1 dependent) apoptosis (**Figure S2D**). Apaf-1 is an ATPase belonging to AP subfamily of STAND ATPase family (Danot *et al*., 2009). The activation of Apaf-1 is executed by ATPase dependent oligomerization with cytochrome c (cyt c) (Riedl and Salvesen, 2007). Unlike NLRC4 whose disk-shaped oligomer is initiated by one ligand protein flagellin binding to one receptor protein NAIP5 which acts as the seed to promote the following one-by-one recruitment of NLRC4, Apaf-1 complex (apoptosome) is initiated by cyt c binding to Apaf-1 in their monomer form and activating ATPase in Apaf-1, which drives the incorporation of cyt c and Apaf-1 together into a complex and constitutes a heptagonal structure with a 1:1 ratio of cyt c to Apaf-1 (Danot *et al*., 2009; Riedl and Salvesen, 2007). Given that NLRC4 is incorporated into the complex alone, a catalyzer is required for its ATPase activation otherwise the complex cannot form in the right structure. Taken together, the oligomeric assembly of STAND ATPase family members requires their ATPase activation either by their binding partner/ligand or a catalyzer, and we propose that our catalyzer model should be applicable to the ATPase dependent assembly of all oligomeric complexes of which the main structural constituent is progressively incorporated into the complex one by one. The catalyzer dependent and independent assembly of signaling complexes may represent two categories of ligand induced ATP hydrolysis (energy)-dependent signaling complex formation (**Figure S7J**).

Potentiation of nucleoside-triphosphatase (NTPase) activity may be a common property of PELO since it increases the GTPase activity of its partner HBS1L in the ribosome rescue pathway (Chen et al., 2010; Graille et al., 2008; Pisareva et al., 2011; Shoemaker et al., 2010). This newly emerged function of PELO in NLRs activation does not utilize the machinery of ribosome rescue. But ribosome rescue and inflammasomal NLR activation can compete for the use of PELO, which could be the way by which PELO limits the activation of NLRs. The common requirement of PELO by all NLRs for their activation should be a key regulatory mechanism in NLR-mediated immune responses, and the shared usage of PELO might create a network link between immune responses and protein translational control machinery.

In the *in vivo* studies, PELO was reported to be involved in several biological processes (Castrillon *et al*., 1993; Liakath-Ali *et al*., 2018; Terrey *et al*., 2021; Wu *et al*., 2014; Wu *et al*., 2018) such as the reproduction of animals. Male *Pelo*-null mutants of *Drosophila* exhibited spermatogenic arrest at the G2/M boundary of the first meiotic division (Eberhart and Wasserman, 1995), and PELO deficient females also showed impaired fertility resulting from a disrupted balance between the self-renewal and differentiation of germline stem cells (GSCs) (Xi et al., 2005). Conditional depletion of *Pelo* in postnatal mice causes progressive spermatogonial stem cell loss and sterility, though later spermatogenesis appears normal before germline exhaustion (Raju et al., 2015). Several non-PRR NLRs exhibit highly restricted expression in mammalian germline and appear to be involved in developmental processes such as spermatogenesis. A study that analyzed the expression of NLRP genes in humans and rhesus macaques (*Macaca mulatta*) also came to the conclusion that the majority of NLRP genes are expressed in the gametes and early embryos of primates (Tian et al., 2009). These observations could be hints for the functional linkage between certain NLRs and PELO in many physiological processes.

It is intriguing to discover that a component of translational quality control machinery also functions *via* a different mechanism in innate immune responses. Whether PELO could be utilized for designing drugs for the therapeutic modulation of inflammatory diseases linked to NLRs activation awaits further investigation. It would also be an interesting topic in the future to understand the evolutionary convergence and divergence of NLRs-related innate immune system and PELO-related ribosome-associated quality control machinery.

## ACKNOWLEDGEMENTS

We thank Chuan-Qi Zhong and Yuting You for mass spectrometry data analysis, Yifei Liu for help with histology, Jiongcong Lu, Bo Liang and Rangxin Peng for help with mouse model, Zhonghan Li (Sichuan University) for NLRP14 plasmid, Lu Zhou for help with proofreading. This work was supported by the National Natural Science Foundation of China (81788101 to J.H., 32170751 to Z.-H.Y.), National Key R&D Program of China (2020YFA0803500 to J.H.), the CAMS Innovation Fund for Medical Science (2019-I2M-5-062 to J.H.).

## AUTHOR CONTRIBUTIONS

X.W., Z.-H.Y. and J.H. designed the research, performed data analyses and wrote the manuscript. X.W. and Z.-H.Y. performed the experiments. J.W. generated and provided relevant mice. J.H. conceived and supervised the study.

---

**Figure S1.**
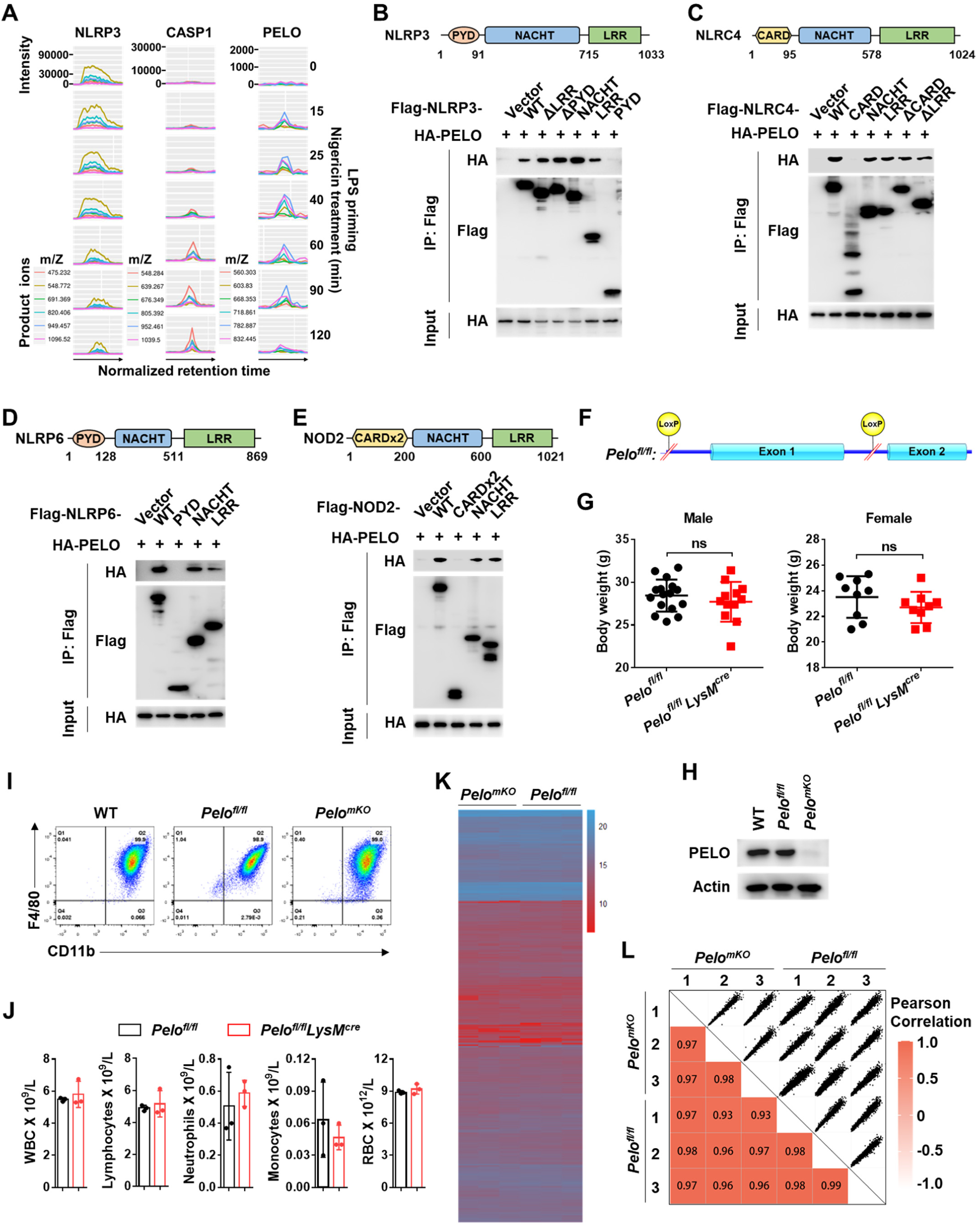
Interaction of PELO with NLRP3 and other NLRs, interacting domain mapping, and generation and characterization of *Pelo* gene knockout mice, related to Figure 1. (**A**) NLRP3 complex was affinity-purified from Flag-NLRP3 reconstituted J774 cells after LPS (100 ng/ml) priming for 4 hours and then nigericin (10 µM) stimulation for 0, 5, 15, 40, 60, 90 or 120 min, digested with trypsin and subjected to high-sensitive quantitative MS analysis. MS data were analyzed with Group-DIA and peptide intensities were extracted. The extracted ion chromatogram (XIC) peaks of multiple product ions of one representative peptide for each of these proteins were shown. Note: KVQTESSTGSVGSNRVR is the sequence of PELO peptide detected. (**B**-**E**) Flag-tagged full length (WT) or truncated NLRP3 (**B**), NLRC4 (**C**), NLRP6 (**D**) or NOD2 (**E**) was co-expressed with HA-tagged PELO in HEK293T cells. The cell lysates were immunoprecipitated with anti-Flag antibodies, then the cell lysates and immunoprecipitates were analyzed by immunoblotting with antibodies as indicated. Schematics of mouse NLRP3, NLRC4, NLRP6 and NOD2 domain architecture were shown. (**F**) Genomic locations of *loxP* sites in the *Pelo* locus of *Pelo^fl/fl^* mouse strain generated. (**G**) Body weights of 18∼20 weeks old mice of indicated genotypes (n ≥ 9 mice per genotype). ns, not significant. (**H**) Immunoblot analysis of the levels of PELO protein in BMDMs. (**I**) Flow cytometry analysis of *ex vivo* differentiated BMDMs stained for CD11b and F4/80. (**J**) Quantification of the numbers of different immune-cell types from the blood of the indicated mouse strains. WBC, white blood cells; RBC, red blood cells. n = 3 mice per genotype. Data are represented as mean ± SD (**G** and **J**). (**K**) Heat map showing MS analyses of global protein expression in *Pelo^fl/fl^* and *Pelo^mKO^*BMDMs. n = 3 mice per genotype. (**L**) Correlation analysis of protein intensities between any two *Pelo^fl/fl^* and *Pelo^mKO^* BMDMs. The matrix of correlation plots is shown, and the colors represent the indicated Pearson correlation coefficient. All results are representative of at least two independent experiments. See also **Table S1**.

**Figure S2.**
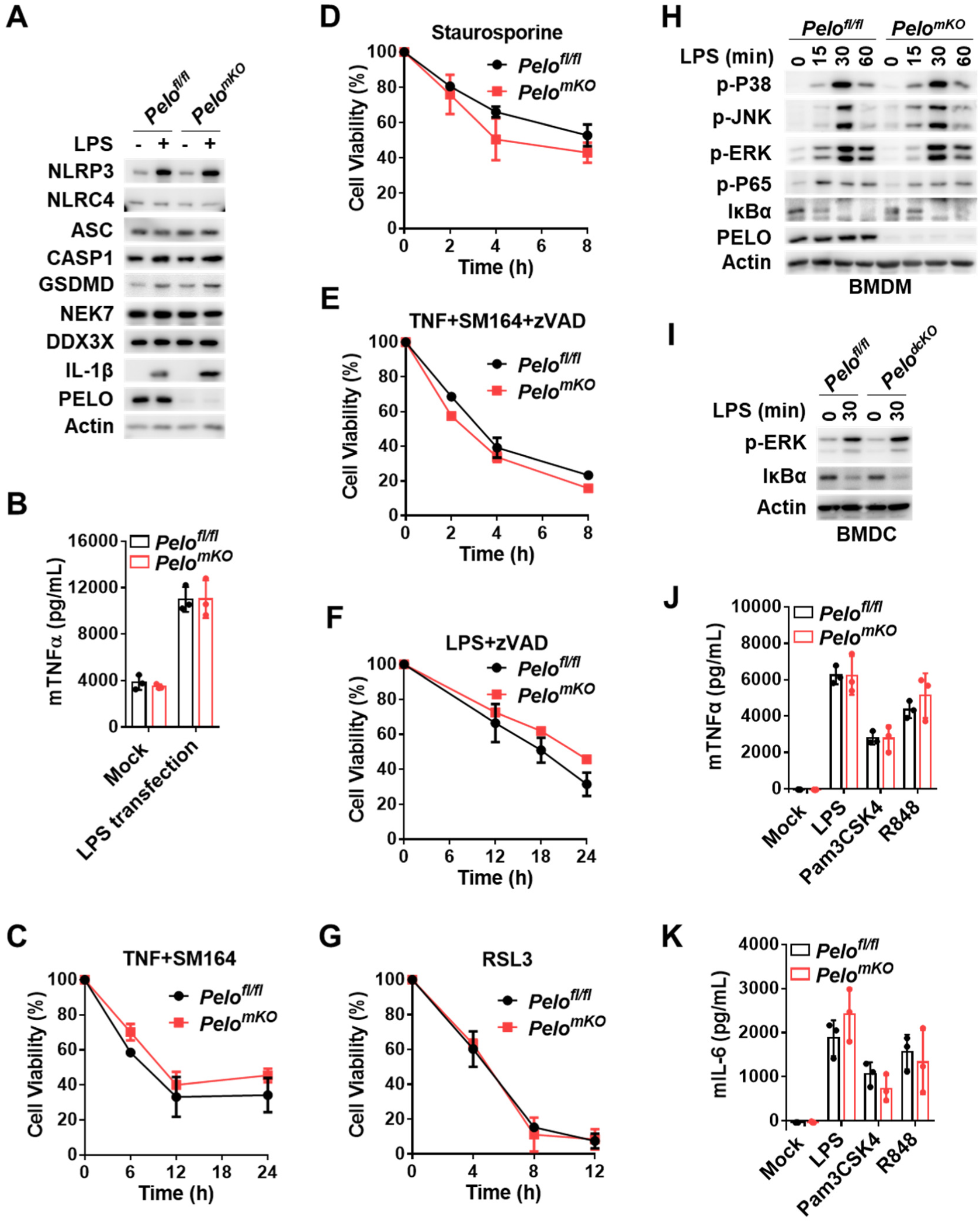
PELO is dispensable for TLRs signaling and the induction of apoptosis, necroptosis and ferroptosis in BMDMs or BMDCs, related to Figure 1. (**A**) *Pelo^fl/fl^* and *Pelo^mKO^* BMDMs were left untreated or stimulated with LPS for 4 hours, and the cell lysates were immunoblotted with indicated antibodies. (B) *Pelo^fl/fl^* and *Pelo^mKO^*BMDMs were primed with Pam3CSK4 (1 μg/ml) for 6 hours and then transfected with LPS (2 μg/ml). 12 hours post-transfection, TNFα secretion from BMDMs was analyzed. n = 3 mice per genotype. (**C-G)** Real-time analysis of cell viabilities in *Pelo^fl/fl^* and *Pelo^mKO^* BMDMs after being stimulated with TNF+SM164 (**C**), Staurosporin (**D**), TNF+SM164+zVAD (**E**), LPS+zVAD (**F**) or RSL3 (**G**). n = 3 mice per genotype. (**H** and **I**) Immunoblot analysis of the cell lysates from *Pelo^fl/fl^* and *Pelo^mKO^*BMDMs (**H**), *Pelo^fl/fl^* and *Pelo^dcKO^* BMDCs (**I**) treated with LPS at indicated time points. (**J** and **K**) TNFα (**J**) and IL-6 (**K**) secretion from *Pelo^fl/fl^* and *Pelo^mKO^* BMDMs treated with the indicated stimuli. n=3 mice per genotype. Data are represented as mean ± SD (**B-G**, **J** and **K**). All results are representative of at least two independent experiments.

**Figure S3.**
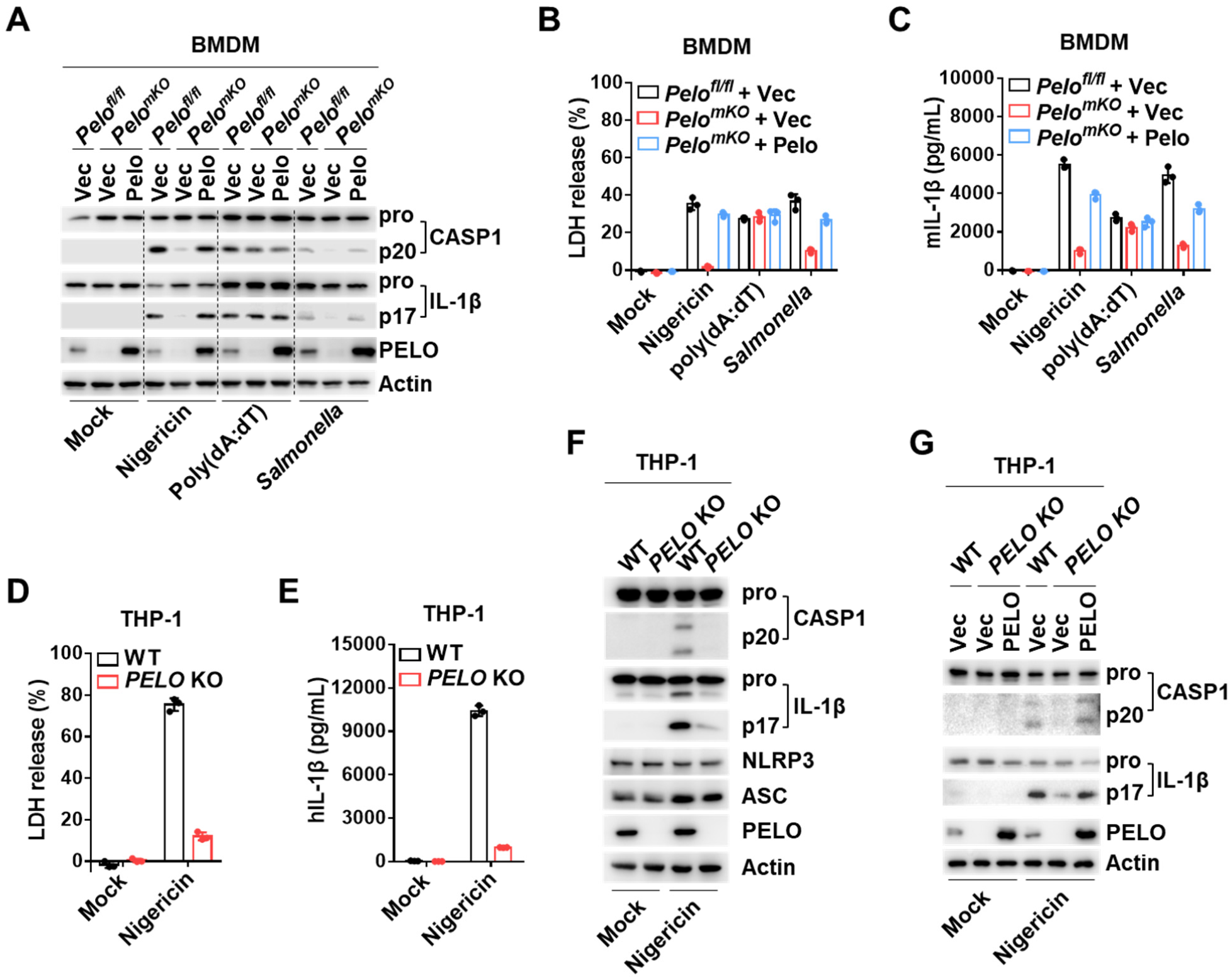
PELO is required for the NLRP3 and NLRC4 inflammasome activation in macrophages, related to Figure 1. (**A**) *Pelo^fl/fl^* BMDMs and the *Pelo^mKO^* BMDMs infected with lentivirus encoding nothing (Vec) or Pelo were primed with LPS and then treated with indicated stimuli. The processed caspase-1 (CASP1) and IL-1β in the pooled cell extracts and supernatants were analyzed by immunoblot. (**B** and **C**) BMDMs were treated as in (**A**). LDH (**B**) and IL-1β (**C**) in culture supernatants were analyzed. Data are represented as mean ± SD of triplicate wells (**B** and **C**). (**D** and **E**) LDH (**D**) and IL-1β (**E**) in culture supernatants of WT and *PELO* KO THP-1 cells that were differentiated by PMA, primed with LPS for 4 hours and then treated with nigericin for 1 hour. Data are represented as mean ± SD of triplicate wells. (**F**) Immunoblot analysis of the processed caspase-1 (CASP1) and IL-1β in the pooled cell extracts and supernatants from WT and *PELO* KO THP-1 cells that were treated as in (**D**). (**G**) WT and *PELO* KO THP-1 cells were reconstituted with Vector (Vec) or PELO expression and treated as in (**D**). The processed caspase-1 (CASP1) and IL-1β in the pooled cell extracts and supernatants were analyzed by immunoblotting. All results are representative of three independent experiments.

**Figure S4.**
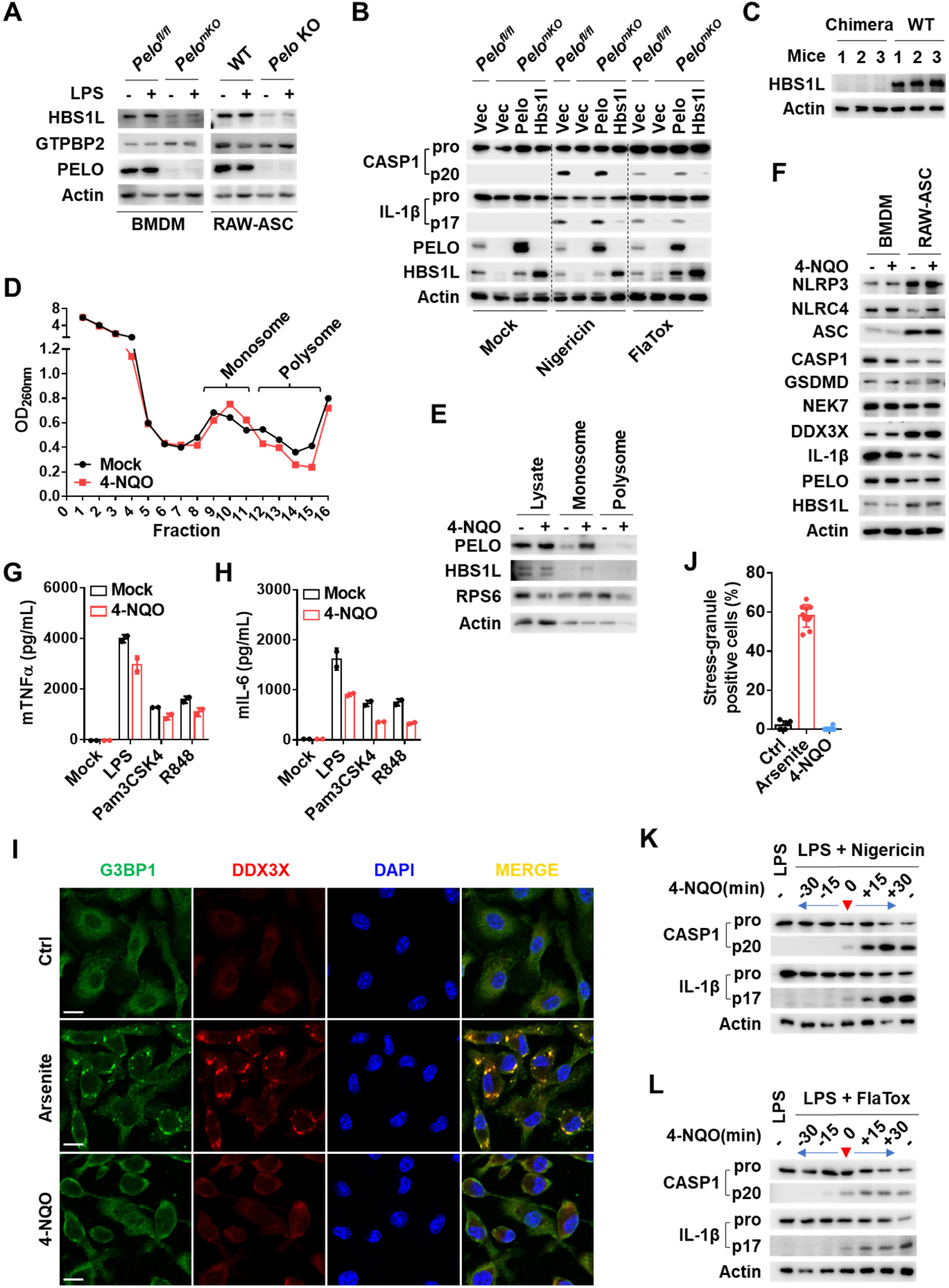
HBS1L and GTPBP2 are not involved in the activation of NLRP3 and NLRC4 inflammasome, and induction of ribosome stalling and activation of NLRP3 or NLRC4 inflammasome antagonize each other, related with Figure 3. (**A**) *Pelo^fl/fl^* and *Pelo^mKO^* BMDMs, wild-type (WT) and *Pelo* KO RAW-ASC cells were left untreated or stimulated with LPS for 4 hours, and the cell lysates were immunoblotted with indicated antibodies. (**B**) *Pelo^fl/fl^* and *Pelo^mKO^* BMDMs infected with lentivirus encoding nothing (Vec), Pelo or Hbs1l were primed with LPS and then treated as indicated. The processed caspase-1 (CASP1) and IL-1β in the pooled cell extracts and supernatants were analyzed by immunoblotting. (**C**) Immunoblot analysis of the levels of HBS1L protein in BMDMs from *Hbs1l^-/-^* chimeric mice and WT mice. (**D** and **E**) Polysome profiling in BMDMs treated with 4-NQO for 1 hour (**D**), and the polysome and monosome fractions were analyzed by immunoblotting with indicated antibodies (**E**). (**F**) LPS-primed BMDMs and RAW-ASC cells were left untreated or stimulated with 4-NQO for 1 hour, and the cell lysates were immunoblotted with indicated antibodies. (**G** and **H**) BMDMs were pre-treated with LPS, Pam3CSK4 or R848 for 4 hours and then treated with 4-NQO for 1 hour. TNFα (**G**) and IL-6 (**H**) secretion from BMDMs were measured. Data are represented as mean ± SD of triplicate wells. (**I**) Representative images for immunofluorescence staining of G3BP1 (green), DDX3X (red) and nuclei (blue) in BMDMs 30 min after treatment with arsenite or 4-NQO. Scale bars represent 10 μm. (**J**) Quantification of the percentage of stress-granule-positive cells after treatment with arsenite or 4-NQO. Data are represented as mean ± SD (n ≥ 6 biologically independent fields of cells). (**K** and **L**) Immunoblot analysis of CASP1 and IL-1β cleavage in the pooled cell extracts and supernatants from LPS-primed BMDMs to which 4-NQO was added 30 and 15 min before, simultaneously with, or 15 and 30 min after the addition of nigericin (**K**) or FlaTox (**L**). All results are representative of three independent experiments. See also **Table S2**.

**Figure S5.**
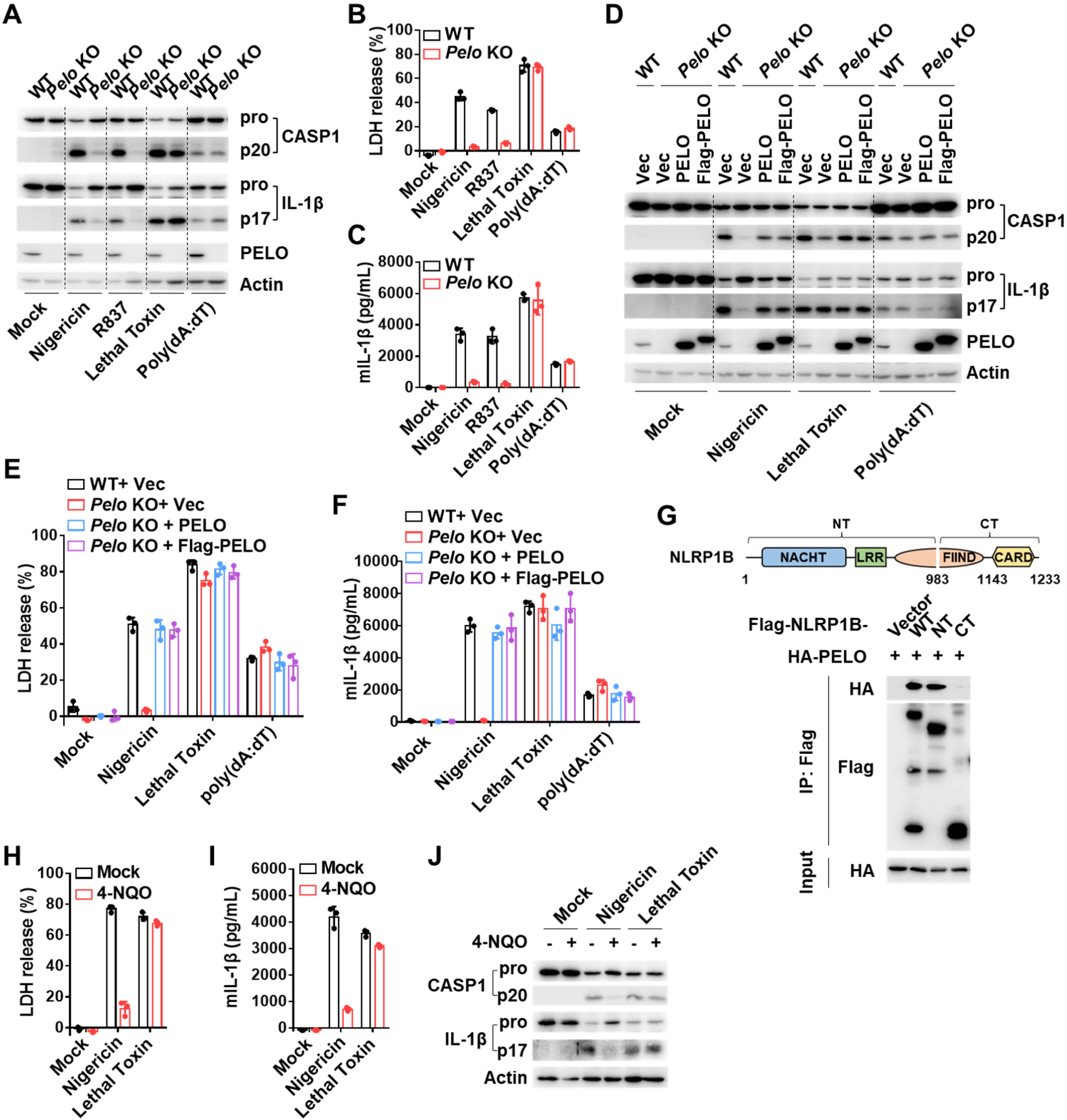
PELO does not participate in lethal toxin induced NLRP1B activation and 4-NQO has no effect on PELO-independent activation of NLRP1B inflammasome, related to Figure 3. (**A**) Immunoblot analysis of the processed caspase-1 (CASP1) and IL-1β in the pooled cell extracts and supernatants from RAW-ASC cells that were primed with LPS and then treated with the indicated stimuli. Mock represents cells primed with LPS without further stimulation. (**B** and **C**) RAW-ASC cells were treated as in (**A**). LDH (**B**) and IL-1β (**C**) in culture supernatants were analyzed. (**D**) WT and *Pelo* KO RAW-ASC cells infected with lentivirus encoding nothing (Vec), PELO or Flag-PELO were primed with LPS and then treated with indicated stimuli. The processed caspase-1 (CASP1) and IL-1β in the pooled cell extracts and supernatants were analyzed by immunoblotting. (**E** and **F**) RAW-ASC cells were treated as in (**D**). LDH (**E**) and IL-1β (**F**) in culture supernatants were analyzed. (**G**) Flag-tagged WT or truncated NLRP1B was co-expressed with HA-tagged PELO in HEK293T cells. The cell lysates were immunoprecipitated with anti-Flag antibodies, and then the cell lysates and immunoprecipitates were analyzed by immunoblotting as indicated. Schematics of mouse NLRP1B domain architecture was shown. (**H-J)** LPS-primed RAW-ASC cells were left untreated or pre-treated with 4-NQO for 30 min and then treated with the indicated stimuli. LDH (**H**) and IL-1β (**I**) in culture supernatants were measured. The processed CASP1 and IL-1β in the pooled cell extracts and supernatants were analyzed by immunoblotting (**J**). Data are represented as mean ± SD of triplicate wells (**B**, **C**, **E**, **F**, **H** and **I**). All results are representative of three independent experiments.

**Figure S6.**
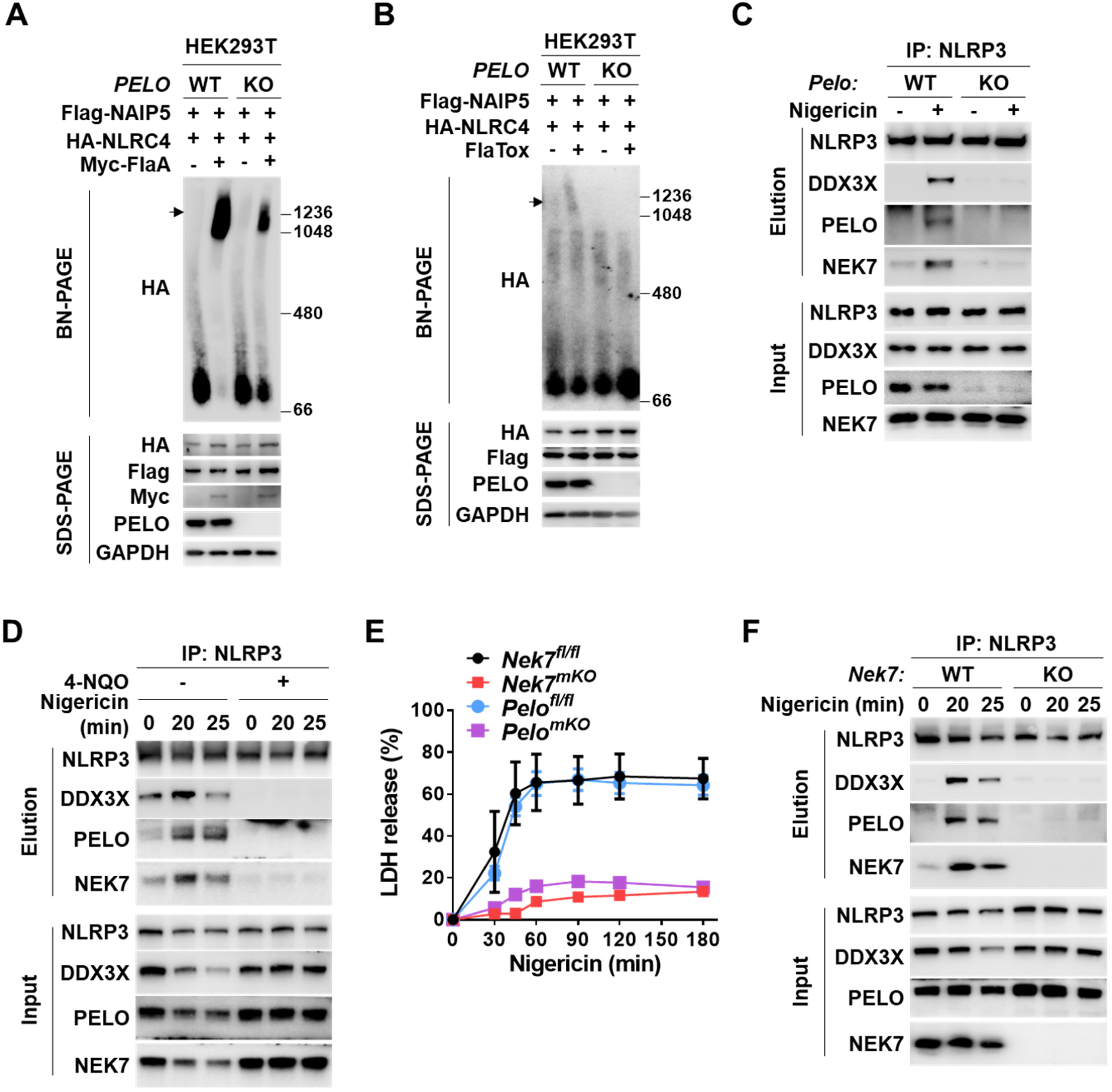
PELO is required for the assembly of NLRP3 and NLRC4 inflammasomes, related to Figure 5. (**A**) WT and *PELO* KO HEK293T cells were transfected with expression vectors of Flag-NAIP5, HA-NLRC4 and Myc-FlaA in different combinations as indicated. 24 hours later, the cell lysates were resolved by blue native PAGE (BN-PAGE) for detection of NLRC4 in high-molecular-weight complexes (arrow), or by SDS-PAGE for detection of expression of each transfected gene (bottom). Immunoblot analyses were performed thereafter. (**B**) WT and *PELO* KO HEK293T cells were transfected with expression vectors of Flag-NAIP5 together with HA-NLRC4. 24 hours later, the cells were treated with FlaTox for 6 hours, and then the cell lysates were analyzed as in (**A**). (**C**) LPS-primed WT and *Pelo^mKO^*BMDMs were stimulated with nigericin as indicated. The cell lysates were immunoprecipitated with anti-NLRP3 antibodies, followed by immunoblot analysis as indicated. **(D)** LPS-primed BMDMs were left untreated or pre-treated with 4-NQO for 30 min and then treated with nigericin. The cell lysates were immunoprecipitated with anti-NLRP3 antibodies, followed by immunoblot analysis as indicated. (**E**) Real-time analysis of LDH release from BMDMs of indicated genotypes that were primed with LPS and then stimulated with nigericin. Data are represented as mean ± SD of triplicate wells. (**F**) Same as in (**C**) except *Nek7^mKO^* BMDMs were used. All results are representative of three independent experiments.

**Figure S7.**
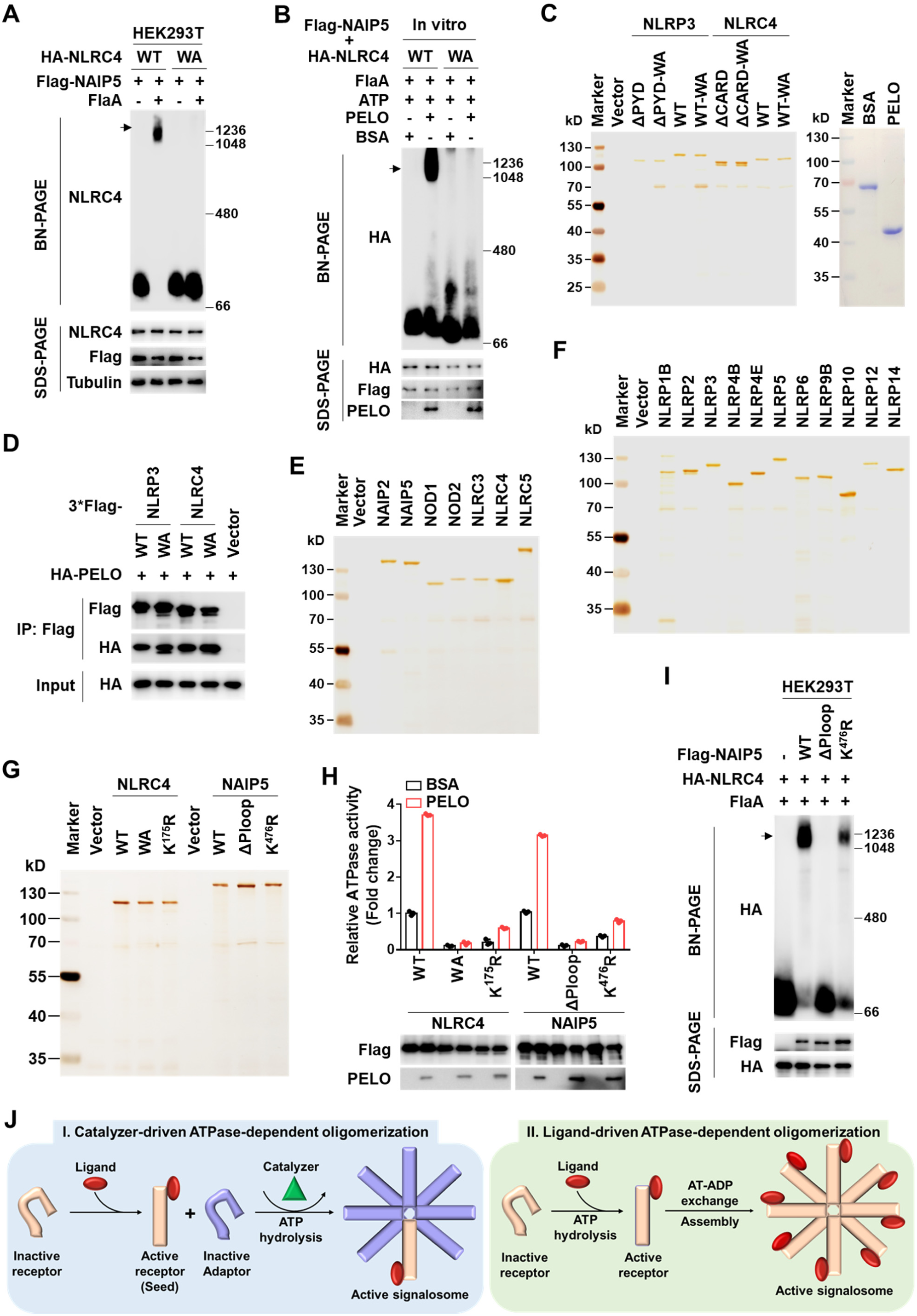
PELO potentiates the ATPase activity of NLRs, related to Figure 6. (**A**) HEK293T cells were transfected with expression vectors of Flag-NAIP5, HA-NLRC4, and Myc-FlaA in different combinations as indicated. 24 hours later, the cell lysates were resolved by blue native PAGE (BN-PAGE) for detection of NLRC4 in high-molecular-weight complexes (arrow), or by SDS-PAGE for detection of expression of each transfected gene (bottom). Immunoblot analyses were performed thereafter. (**B**) NLRC4 and NAIP5 containing *PELO* KO HEK293T cell lysates were incubated with recombinant His-PELO protein as indicated in the presence of flagellin (FlaA) and ATP, and then the reaction mixtures were analyzed by BN-PAGE. (**C**, **E-G**) Representative image of silver staining of the purified Flag-tagged proteins as indicated and Coomassie blue staining of recombinant His-tagged PELO protein. (**D**) Flag-tagged WT or mutant NLRP3 or NLRC4 was co-expressed with HA-tagged PELO in HEK293T cells. The cell lysates were immunoprecipitated with anti-Flag antibodies, and then the cell lysates and immunoprecipitates were analyzed by immunoblotting as indicated. (**H**) ATPase activities of purified Flag-WT or mutant NLRC4 and NAIP5 in the presence of BSA or recombinant His-PELO protein. Folds of change were shown. Data are represented as mean ± SD of triplicates. (**I**) Flag-tagged WT or mutant NAIP5 was transfected into HEK293T cells with WT NLRC4 and FlaA and the cell lysates were analyzed as in (**A**). (**J**) A schematic diagram for the two categories of ATPase-dependent oligomeric signaling complex formation. All results are representative of at least two independent experiments.

## STAR★METHODS

Detailed methods are provided in the online version of this paper and include the following:

### KEY RESOURCES TABLE

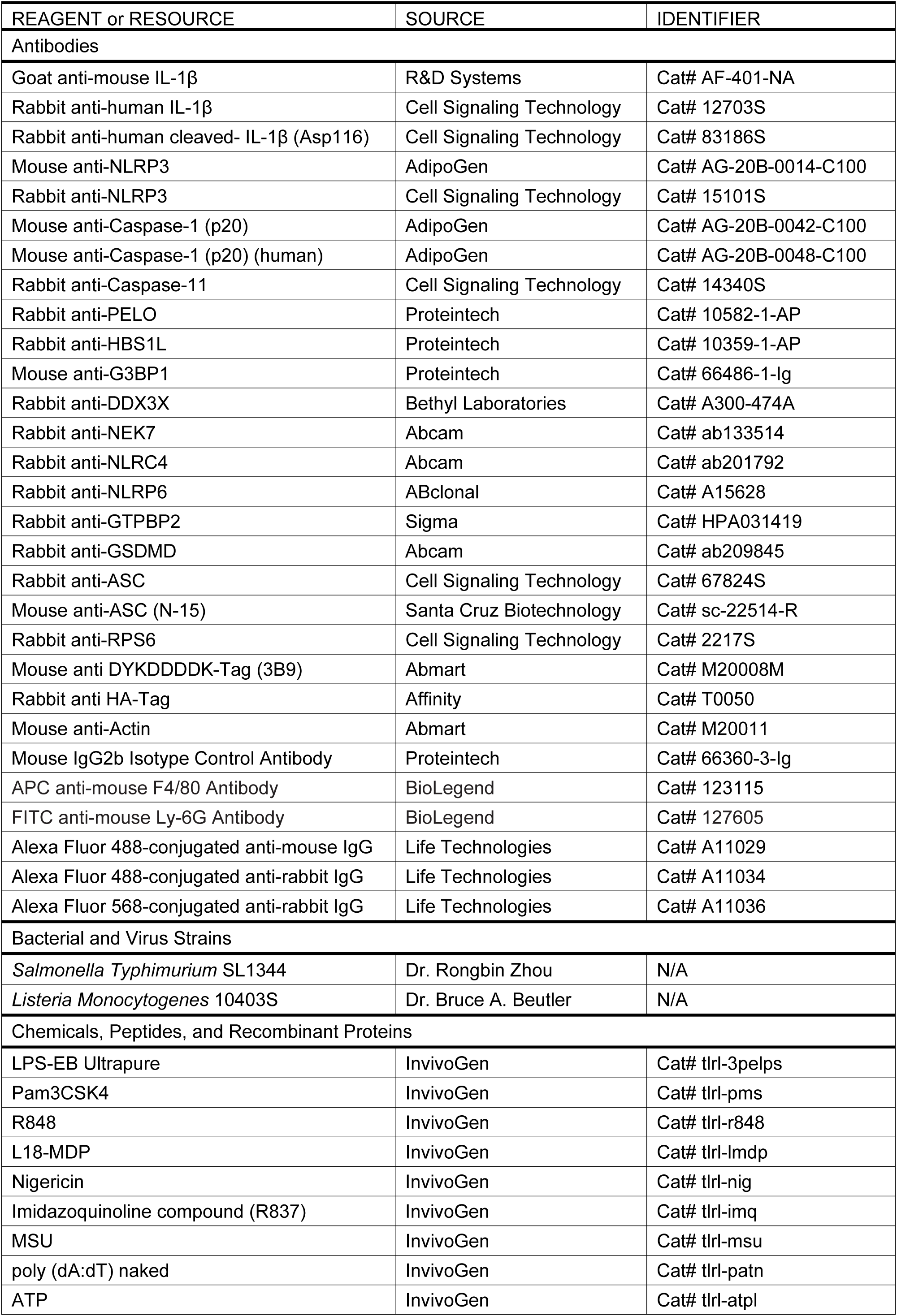
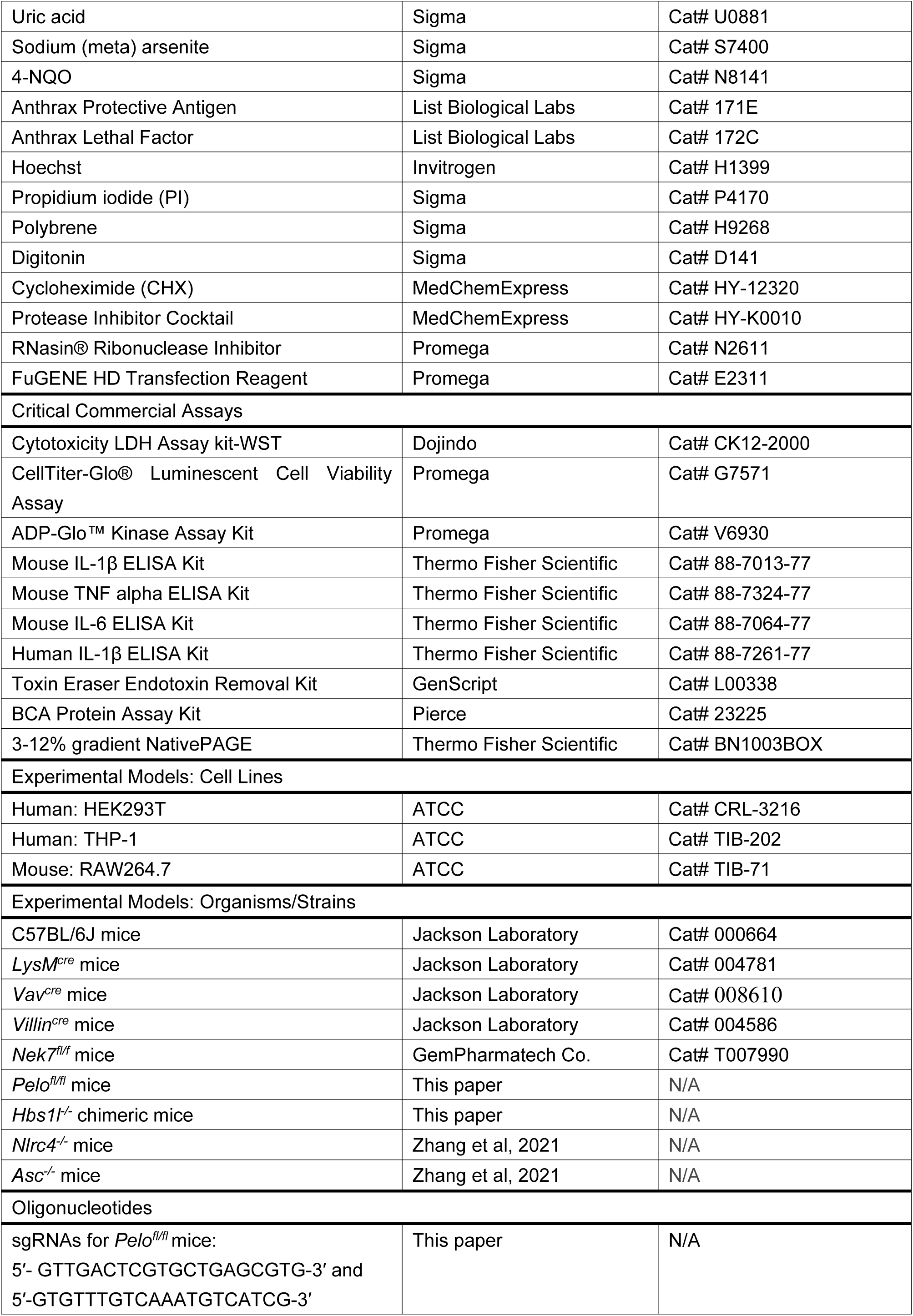
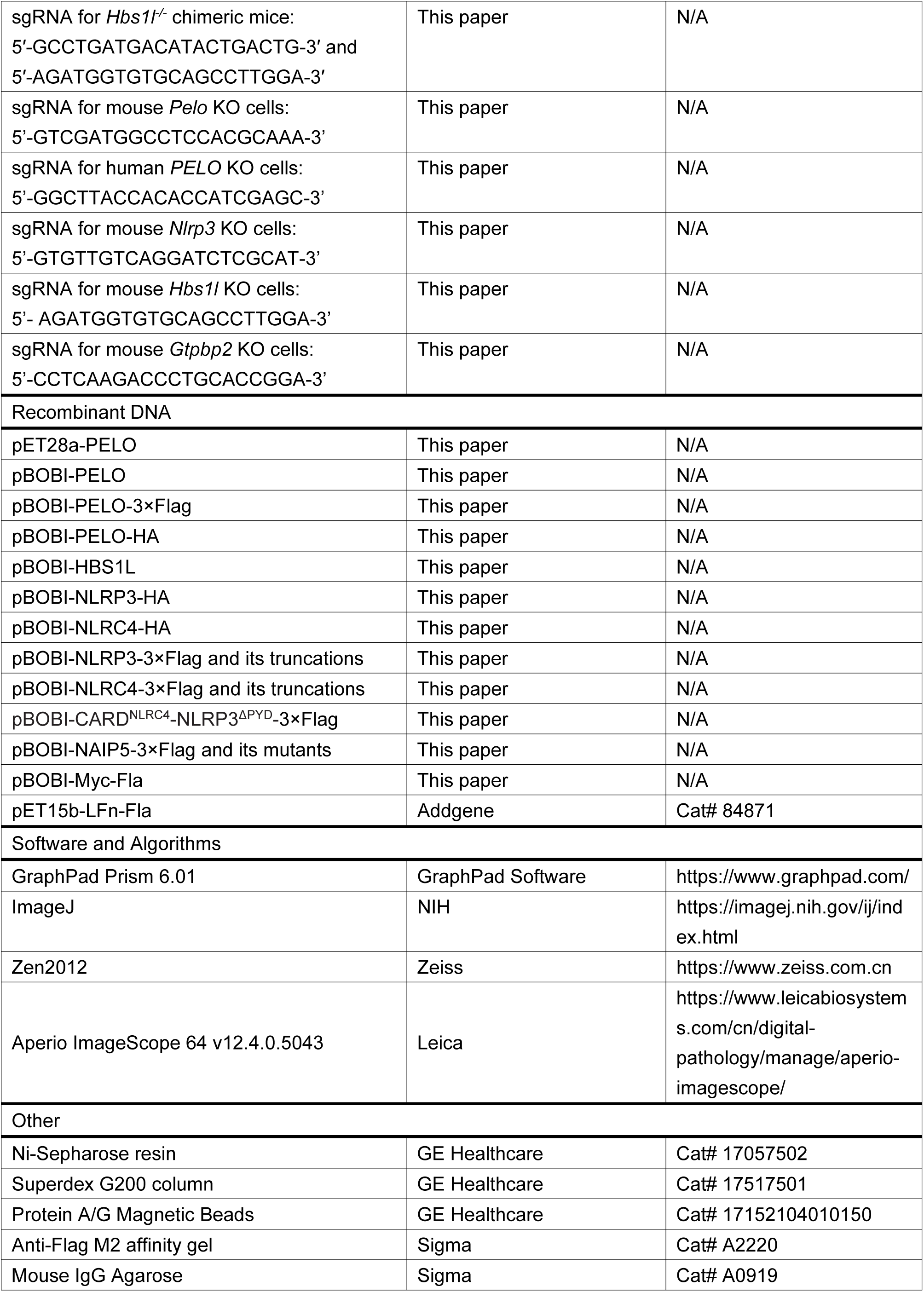

### RESOURCE AVAILABILTY

#### Lead contact

Further information and requests for resources and reagents should be directed to the lead contact, Jiahuai Han.

#### Materials availability

All plasmids, reagents, cell lines and mouse lines generated in this study are available from the lead contact.

#### Data and code availability

This study did not generate datasets/code.

### EXPERIMENTAL MODEL AND SUBJECT DETAILS

#### Cell culture

BMDMs were generated by differentiating bone marrow progenitors from the tibia and femur for 7 days in Dulbecco’s modified Eagle’s medium (DMEM) supplemented with 10% (v/v) fetal bovine serum (FBS) and 30% (v/v) L929-conditioned medium. BMDCs were generated by differentiating bone marrow progenitors from the tibia and femur of *Pelo^fl/fl^* and *Pelo^fl/fl^Vav^cre^* mice for 8 days in RPMI 1640 containing 10% (v/v) heat-inactivated FBS, 5×10^-5^ M β-Me and 20 ng ml^-1^ GM-CSF. RAW-ASC cells were generated as described (He *et al*., 2015). RAW-ASC, RAW264.7 and HEK293T cells were grown in DMEM supplemented with 10% FBS. TPH-1 cells were cultured in RPMI 1640 medium supplemented with 10% FBS. All cells were grown at 37 °C in a 5% CO2 incubator.

#### Mice

*LysM^cre^* (Jax stock, 004781) mice, *Vav^cre^* (Jax stock, 008610) mice and *Villin^cre^* mice (Jax stock, 004586) were purchased from the Jackson Lab. *Nek7^fl/fl^* mice were purchased from the GemPharmatech Co., Ltd. *Pelo^fl/fl^* mice were generated by a previously described androgenetic haploid embryonic stem cells (AG-haESCs)-based method (Zhong et al., 2015). Briefly, the high-efficient sgRNAs targeting *Pelo* were: 5′-GTTGACTCGTGCTGAGCGTG-3′ and 5′-GTGTTTGTCAAATGTCATCG-3′, which were cloned into a modified pSpCas9 (BB)-P2A-mcherry-puromycin-sgRNA vector. The homologous-recombination template plasmid was constructed by insertion of a 1,087-bp genomic fragment, along with a 1-kb genomic fragment as 5′-homology-arm and 0.9-kb genomic fragment as 3′-homology-arm. Constructed sgRNA and template plasmids were transferred into cultured AG-haESCs. 20 hours after transfection, mCherry-positive haploid cells were sorted with FACSAria™ III flow cytometer (BD) and re-cultured at a density of 7,000 cells per well (6-well plate). Each single clone was genotyped by genomic fragment sequencing, and those corrected clones were retained. The picked clones were then digested into single cells, and intra-cytoplasmically injected to the mature oocyte from the C57BL/6J × DBA2 F1 generation mice. After being cultured for 24 hours, 2-cell ESCs were transplanted into prepared pseudopregnant ICR female mice and the offspring were further confirmed by genomic fragment sequencing. The floxed-offspring mice were back-crossed with C57BL/6J wild-type mice for at least 6 generations for further studies. *Hbs1l^-/-^* chimeric mice were generated by co-microinjection of *in vitro* transcribed Cas9 mRNA and sgRNAs into the C57BL/6J zygotes. The target sequence in the sgRNA vector was 5′-AGATGGTGTGCAGCCTTGGA-3′ for mouse *Hbs1l*. Consistent with a recent report (Terrey *et al*., 2021), whole-body genetic knockout of *Hbs1l* in mice is lethal. We obtained three *Hbs1l^-/-^* chimeric mice of which the two alleles of *Hbs1l* in bone marrow cells were disrupted. *Nlrc4^−/−^* mice, *Asc^−/−^* mice and *Casp1^−/−^* mice were used as previously described (Zhang *et al*., 2021). Sample sizes for mouse experiments were empirically determined, and mice were randomly assigned to the control or experimental groups. For all experiments, age- and sex-matched 6 to12 week-old littermates were used as indicated in each figure. All mice were in the C57BL/6J background and housed in a conventional environment under a 12-hour light:dark cycle at Xiamen University Laboratory Animal Center. All mouse experiments were approved by the Institutional Animal Care and Use Committee and were in strict accordance with good animal practices as defined by the Xiamen University Laboratory Animal Center.

### METHOD DETAILS

#### Generation of knockout cell lines

The target sequences in the gRNA vector were 5’-GTCGATGGCCTCCACGCAAA-3’ for mouse *Pelo,* 5’-GGCTTACCACACCATCGAGC-3’ for human *PELO*, 5’-GTGTTGTCAGGATCTCGCAT-3’ for mouse *Nlrp3*, 5’-AGATGGTGTGCAGCCTTGGA-3’ for mouse *Hbs1l* and 5’-CCTCAAGACCCTGCACCGGA-3’ for mouse *Gtpbp2*. To construct the knockout cell lines, gRNA was transduced into indicated cell line by lentiviral delivery. Cells were then subjected to blasticidin selection. Single-cell clones were isolated from the selected pool by limiting dilution cloning in 96-well plates and then screened for indicated gene expression by western blot. Selected knockout clones were verified by DNA sequencing.

#### Lentivirus production and infection

The lentiviral vectors carrying cDNAs or gRNAs of interest were transfected into 293T cells in the presence of lentivirus-packing plasmids (PMDL/REV/VSVG) by the calcium phosphate precipitation method. 12 hours later, cell culture medium was changed and the virus-containing medium was collected 30 hours later. Primary BMDMs were infected with lentiviral particles in the presence of 8 μg ml^−1^ polybrene and then centrifuged at 2,500 rpm for 60 min on day 3 of the differentiation protocol. For other cells, virus containing medium in the presence of 10 μg ml^-1^ polybrene was added to cells plated, followed by centrifugation at 2,500 rpm for 30 min.

#### Recombinant protein preparation

pET15b LFn-Fla (Addgene plasmid #84871) was a gift from Dr. Russell Vance. The LFn-Fla was expressed and purified as described (von Moltke *et al*., 2012). The endotoxin was removed with Toxin Eraser Endotoxin Removal Kit (L00338, GenScript). The C-terminal 6×His-tagged PELO protein was expressed in *E. coli* BL21 (DE3) strain (Novagen, Merck) by overnight culturing at 30 °C in an auto-induction medium. The PELO protein was first purified with Ni^2+^-NTA-agarose (Qiagen) and further purified by Superdex 200 10/30 prepacked column (GE Healthcare). The protein concentration was determined using the BCA method with BSA as the standard.

#### NLR proteins preparation

To produce purified NLR proteins, HEK293T cells were transiently transfected with plasmids encoding 3×Flag-tagged NLRs. Plasmid encoding nothing (vector) was used as control. 36 hours after transfection, the cells were washed twice in cold PBS and lysed in lysis buffer (50 mM HEPES, pH7.4, 150 mM NaCl, 1% NP-40) supplemented with Protease Inhibitor Cocktail. The lysates were incubated at 4℃ on a rotation platform for 30 min and then centrifuged at 20, 000 g for 30 min at 4℃. The supernatants were incubated with IgG-Agarose at 4℃ on rotation for 2 hours. The pre-cleaned lysates were incubated with Anti-Flag M2 Affinity Gel for 3 hours at 4℃ on rotation, and then washed with wash buffer A (50 mM HEPES, pH7.4, 300 mM NaCl, 1% NP-40) for three times, followed by another three times of wash with wash buffer B (50 mM HEPES, pH7.4, 150 mM NaCl, 5% Glycerol, 10 mM MgCl_2_, 0.1% NP-40). The NLR proteins were finally eluted with wash buffer B containing 200 μg ml-1 3×Flag peptide.

#### ATPase activity assay

Assay was carried out using the ADP-Glo kinase assay kit in 384 Flat White Plates (142761, Thermo Fisher Scientific). Respective purified NLR proteins were incubated at 37℃ with BSA or PELO-His protein for 30 min in the reaction buffer (50 mM HEPES, pH7.4, 150 mM NaCl, 5% Glycerol, 10 mM MgCl_2_, 0.1% NP-40, 1 mM DTT). ATP (20 μM in final) was then added, and the mixture was further incubated at 37℃ for another 40 min. The reaction is stopped by addition of ADP-Glo reagent and further incubated for 40 min at room temperature. After addition of kinase detection buffer, samples were incubated for another 40 min and luminescence read out using a Spark 20M microplate reader (Tecan) with an integration time of 1 s per well.

#### Inflammasome activation

Macrophages were plated in 96-well plates at 1×10^5^ cells per well. Unless indicated, mice macrophages were primed with 100 ng ml^-1^ ultrapure LPS for 4 hours, followed by stimulation with 5 mM ATP (30 min), 5 μM nigericin (1 hour), 20 μg ml^-1^ R837 (1 hour), 200 μg ml^-1^ MSU (6 hours), 2 μg ml^-1^ poly(dA:dT) (2 hours), 2 μg ml^-1^ LFn-Flagellin together with 2 μg ml^-1^ PA proteins (1 hour), *Salmonella* (multiplicity of infection (m.o.i)=10, 1 hour), 2 μg ml^-1^ Lethal factor together with 2 μg ml^-1^ PA proteins (90 min). THP-1 cells were primed with 1 mg ml^-1^ LPS for 4 hours, followed by stimulation with 5 μM nigericin (1 hour). For non-canonical inflammasome activation, BMDMs were primed with 1 μg ml^-1^ Pam3CSK4 for 6 hours and then were transfected with 2 μg ml^-1^ LPS by using 0.25% v/v FuGENE HD (Promega). After stimulation, culture supernatants and cell lysates were collected together for immunoblotting analysis and the cell supernatants were used for LDH assay and ELISA analysis.

#### Cytotoxicity assay

Relevant cells were treated as indicated. Pyroptotic cell death was measured by the LDH assay using Cytotoxicity LDH Assay kit-WST (Dojindo Molecular Technologies). Cell viability was measured by the CellTiter-Glo Luminescent Cell Viability Assay (Promega).

#### ELISA

The supernatants from cells or serum from mice were assayed for mouse IL-1β, IL-6, TNFα (Thermo Fisher Scientific) according to the manufacturer’s instructions.

#### Stimulation with endotoxin *in vivo*

Mice were injected intraperitoneally with 10 mg kg^−1^ LPS (Escherichia coli 0111: B4; Sigma). The blood samples were collected 3 hours later. Serum cytokines were measured by ELISA.

#### Infection *in vivo*

Lethal *L. monocytogenes* (10403S) infection was established by infecting mice with 1×10^6^ c.f.u. bacteria administered intraperitoneally (i.p.) in 200 μl PBS. For *S. typhimurium* (SL1344) infection, mice were injected intraperitoneally with 1×10^3^ c.f.u. bacteria in 200 μl PBS. The survival rate of the mice was checked every day.

#### MSU-induced mouse peritonitis

MSU crystals were prepared as previously described (Schiltz et al., 2002). Mice were injected intraperitoneally with 1 mg MSU dissolved in 0.5 ml sterile PBS. After 3 hours, the mice were euthanized and peritoneal cavities were flushed with 2 ml cold PBS. PECs were collected and stained with APC anti-mouse F4/80 Antibody (123115, BioLegend) and FITC anti-mouse Ly-6G Antibody (127605, BioLegend). Flow cytometry data were acquired using a Fortessa X-20 flow cytometer (BD) and analyzed with FlowJo software (FlowJo and Illumina). IL-1β was measured by ELISA.

#### FlaTox injection

Mice were injected intravenously (tail vein) with 0.2 μg/g body weight of LFn-Fla combined with 0.2 μg/g PA. Rectal temperature was determined using a MicroTherma 2T Hand Held Thermometer (Braintree scientific).

#### Immunofluorescence staining and imaging

After stimulation, BMDMs were washed twice with PBS followed by fixation for 15 min at room temperature in freshly prepared 4% paraformaldehyde. The fixed cells were then permeabilized in 0.25% Triton X-100/PBS and non-specific binding was blocked with 3% BSA in PBS. Cells were incubated with the following antibodies for 2 hours at room temperature: anti-G3BP1 (Proteintech, 66486-1-Ig; 1:200), anti-DDX3X (Bethyl Laboratories, A300-474A; 1:200). The secondary antibodies used were Alexa Fluor 568-conjugated anti-rabbit IgG (Life Technologies, A11036; 1:500), Alexa Fluor 488-conjugated anti-mouse IgG (Life Technologies, A11029; 1:500). Cells were counterstained with Hoechst to visualize nuclei. All images were acquired on a Zeiss LSM 780 laser scanning confocal microscope using a 40×objective. Unprocessed images were analyzed by ImageJ software.

#### ASC speck staining and ASC oligomerization assay

Macrophages were seeded overnight onto glass bottom dishes and then stimulated with indicated inflammasome stimuli. After stimulation, cells were fixed with 4% paraformaldehyde, permeabilized with 0.1% Triton X-100, and blocked with 3% BSA. Cells were stained with anti-ASC antibody (CST, 67824S; 1:200) and Alexa Fluor 488-conjugated anti-rabbit IgG (Life Technologies, A11034; 1:500). Hoechst was used to stain nuclei. Images were captured using a 20×objective on a Zeiss Axioimager D2 upright microscope and subsequently processed using ImageJ software. For ASC oligomerization assay, cells were lysed with TBS buffer (50 mM Tris-HCl, 150 mM NaCl, pH7.4) containing 0.5% Triton X-100 and EDTA-free protease inhibitor cocktail. The cell lysates were centrifuged at 6,000 g for 15 min at 4°C. Supernatants were transferred to new tubes as Triton-soluble fractions. The Triton-insoluble pellets were washed twice with TBS buffer and then suspended in 200 μl TBS. The resuspended pellets were then cross-linked at room temperature for 30 min with 2 mM disuccinimidyl suberate (Pierce) and then were centrifuged for 15 min at 6,000 g. The pellets were dissolved in SDS sample buffer.

#### Blue native PAGE

Cells were lysed with ice-cold Native-PAGE lysis buffer (50 mM Bis-tris, 50 mM NaCl, 10% (w/v) glycerol, 0.001% Ponceau S, 1% digitonin, 1× EDTA-free protease inhibitor cocktail, pH 7.2) for 30 min. After 20,000 g spin for 30 min, supernatants were equalized after quantification of total protein using the BCA protein assay (Pierce), mixed with 0.25% Coomassie G-250, and then separated by NativePAGE 3-12% Bis-Tris Gel (Thermo Fisher Scientific). Native gels were soaked in 10% SDS solution for 5 min before being transferred to PVDF membranes (Millipore), followed by conventional western blotting.

#### NLRC4 oligomerization assay

HEK293T cells were seeded into 12-well plate 12 hours before transfection with indicated combinations of plasmids, and collected for blue native PAGE analysis 24 hours post transfection. For FlaTox induced NLRC4 oligomerization analysis, HEK293T cells were co-infected with lentivectors expressing NLRC4 and NAIP5. 48 hours after infection, cells were seeded into 24-well plate. 12 hours later, LFn-Flagellin (10 μg ml^-1^) together with PA proteins (10 μg ml^-1^) or Lipo2000 was added into the culture medium and incubated for another 6 hours. The cells were collected and subjected to blue native PAGE analysis. For in vitro NLRC4 inflammasome reconstitution, HA-NLRC4 and Flag-NAIP5 expressing *PELO* KO HEK293T cells were lysed with ice-cold Native-PAGE lysis buffer (50 mM Bis-tris, 50 mM NaCl, 10% (w/v) glycerol, 0.001% Ponceau S, 1% digitonin, 1× EDTA-free protease inhibitor cocktail, pH 7.2) for 30 min. After 20,000 g spin for 30 min, supernatants were equalized after quantification of total protein using the BCA protein assay (Pierce). Next, cell lysates were mixed with or without ATP (0.2 mM) + Mg^2+^ (2 mM), 200 ng flagellin, 100 ng His-PELO or BSA in a total volume of 15 μl. The mixtures were incubated for 1 hour at 37℃ and then centrifuged for 10 min at 20,000 g. The supernatants were separated by NativePAGE 3-12% Bis-Tris Gel (Thermo Fisher Scientific).

#### Co-immunoprecipitation

HEK293T cells or BMDMs were lysed in NP-40 lysis buffer (25 mM HEPES, pH7.4, 150 mM NaCl, 1% NP-40) supplemented with Protease Inhibitor Cocktail. The lysates were incubated at 4℃ on a rocking platform for 30 min and then centrifuged at 20,000 g for 30 min at 4℃. Flag-tagged proteins were immunoprecipitated with Anti-Flag M2 Affinity Gel for 6 hours at 4℃. For the endogenous interaction assay, the NLRP3 antibody or IgG control antibody was incubated with the Protein A/G Magnetic Beads respectively and then the cell lysates were incubated with the indicated antibody-beads conjugate overnight at 4℃ under gentle rotation. Beads containing protein complexes were washed three times with lysis buffer. The immunocomplexes were eluted in sample buffer and then analyzed by western blotting.

#### Fractionation of NLRC4 complex

Cells were homogenized using a Dounce homogenizer in isotonic buffer (10 mM Tris, pH 7.5, 10 mM KCl, 1.5 mM MgCl_2_, 0.25 M sucrose and protease inhibitor cocktail) and then centrifuged at 20, 000 g for 30 min at 4℃. The supernatant was loaded onto a step-gradient of 20%, 30%, 40%, 50% and 60% sucrose in 10 mM Tris at pH 7.5, 10 mM KCl, 1.5mM MgCl_2_ supplemented with protease inhibitor cocktail and centrifuged for 16 hours at 36,000 rpm (SW60Ti swinging-bucket rotor, Beckman). Fractions of 500 μl were collected manually and mixed with 2×SDS sample buffer for western blotting.

#### Histology

Mice were anaesthetized before being killed. Small-intestinal tissues were flushed with ice-cold PBS, coiled into a ‘Swiss roll’ and fixed in 4% PFA for 24 hours at room temperature. The fixed tissues were dehydrated in ethanol, cleared in xylene, and embedded in paraffin blocks. Five-micrometer sections were cut and mounted on adhesion microscope slides (ZSGB-BIO), and then stained with haematoxylin and eosin (H&E) for analyses. Representative images were captured and processed using identical settings in the Leica Aperio Versa 200 at Xiamen University. The investigators were blinded to allocation when the histology experiments were performed.

#### Polysome profiling

BMDMs (1×10^7^) were first primed with LPS for 4 hours, and then treated with 20 μM 4-NQO for 1 hour. Cells were incubated with 100 μg ml^-1^ cycloheximide (CHX) for 10 min and then washed twice with cold PBS containing CHX (100 μg ml^-1^) and scraped. Cells were then lysed in 250 μl lysis buffer (20 mM HEPES, pH 7.4, 5 mM MgCl_2_, 100 mM KCl, 1% Triton X-100, 0.5 mM DTT, 100 μg ml^-1^ CHX, 200 U ml^-1^ RNAsin RNase inhibitor and Protease-Inhibitor Cocktail). Lysates were cleared by centrifugation at 10,000 g for 10 minutes and 200 μl supernatants were layered onto 1,800 μl 10-50% continuous sucrose gradient and centrifuged at 4 °C for 1 hour at 55,000 rpm in a Beckman TLS-55 rotor. Sixteen fractions, 125 μl each, were collected manually from the top of the gradient and the polysome profile was monitored by RNA absorbance at 260 nm. The samples were mixed with 5×SDS sample buffer and analyzed directly by electrophoresis.

#### Mass spectrometry processing and data analysis

NLRP3 complex was purified and analyzed by high-sensitive quantitative mass spectrometry as previously described (He *et al*., 2015). For global protein expression analysis in BMDMs, protein extraction and digestion were performed using SCASP (Gan et al., 2021). Cells were dissolved in 100 μl 1% SDS/100 mM Tris-HCl (pH 8.5)/10 mM Tris (2-carboxyethyl) phosphine hydrochloride (TCEP)/40 mM Chloroacetamide (CAA) and boiled for 10 min. 30 μl 250 mM HP-β-cyclodextrin were added and pipette-mix to homogenize the solution. Trypsin was subsequently added at protein: trypsin ratio of 100:1. Digestion was performed at 37℃ overnight. Peptides were cleanup using SDB-RPS StageTips. Peptides were analyzed by diaPASEF (Meier et al., 2020) on the timsTOF Pro instrument. Liquid chromatography was performed on an ultra-high-pressure nano-flow chromatography system (Elute UHPLC, Bruker Daltonics). The gradient time is 60 min and the total run time is 75 min including washes and equilibration. LC was coupled online to a hybrid TIMS quadrupole time-of-flight mass spectrometer (Bruker timsTOF Pro). For data-independent acquisition, we adopted the isolation scheme of 25 Da×32 windows to cover 400-1200 mz. diaPASEF (.d) files were loaded into DIA-NN (V.1.7.15) (Demichev et al., 2020). The fasta file without decoys was loaded. “FASTA digest for library-free search” and “Deep learning-based spectra and RTs prediction” were enabled. Quantification mode was set to “Robust LC (high accuracy)”. All other settings were left default. The protein intensity matrix was input into Perseus software. Protein copy number per cell was calculated using the “proteomic ruler” plugin as previously described (Wisniewski et al., 2014). Log2-transformation was performed for relative protein abundance, and the heat map was implemented in the R package “pheatmap”. Pearson correlation coefficients between samples were calculated and visualized.

### QUANTIFICATION AND STATISTICAL ANALYSIS

No statistical methods were used to predetermine sample size. GraphPad Prism software was used for data analysis. Data are shown as mean ± standard deviation (SD). The statistical significance of the differences between the two groups was determined by the unpaired two-tailed *t-*test. Differences in compared groups were considered statistically significantly different with *P* values: ns: *P*≥0.0.5; *: *P* < 0.05; **: *P* < 0.01; ***: *P* < 0.001; ****: *P* < 0.0001.

